# The C-terminal SUMOylation-dependent regulation of αKNL2 governs its centromere targeting and interaction with CENH3

**DOI:** 10.1101/2025.08.18.670863

**Authors:** Manikandan Kalidass, Jitka Vaculíková, Jothipriya Ramakrishnan Chandra, Barbora Králová, Venkata Ganesh Jarubula, Sevim D. Kara Öztürk, Dmitri Demidov, Veit Schubert, David Potesil, Jan J. Palecek, Inna Lermontova

**Affiliations:** Leibniz Institute of Plant Genetics and Crop Plant Research (IPK) Gatersleben, Corrensstrasse 3, D-06466 Seeland, Germany; National Centre for Biomolecular Research, Faculty of Science, Masaryk University, Kamenice 5, 62500 Brno, Czech Republic; Department of Agricultural Genetic Engineering, Ayhan Şahenk Faculty of Agricultural Sciences and Technologies, Niğde Ömer Halisdemir University, 51240, Niğde, Türkiye; Central European Institute of Technology (CEITEC), Masaryk University, Brno, Czech Republic

**Keywords:** Cell division, Centromeres, Kinetochore, SUMOylation, Plant development, Protein-protein interactions

## Abstract

The centromere is a specialized domain that facilitates chromosome segregation during mitosis and serves as the site for kinetochore formation. KINETOCHORE NULL2 (αKNL2) is essential for the recognition and loading of the centromeric histone H3 variant, CENH3, to centromeres. A yeast two-hybrid screen for αKNL2 interactors identified components of the SUMOylation pathway. However, the role of αKNL2 SUMOylation in Arabidopsis has not yet been determined. In this study, we demonstrated that the C-terminal part of αKNL2 interacts with SUMO3 and ULP1d, as shown by BiFC and co-immunoprecipitation assays. Bioinformatic and functional analysis identified three SUMOylation and two SUMO-interacting motif (SIM) sites in the C-terminal region of αKNL2, which are critical for growth, fertility, and chromosome alignment. Of the three SUMOylation sites, Lys474 and Lys511 were the most critical for the centromeric localization of αKNL2, underscoring the importance of αKNL2 SUMOylation for its function. Additionally, both in vitro and in vivo assays showed that αKNL2-C undergoes SUMOylation by SUMO1 or SUMO3. The SUMO protease mutant, *ulp1d-2* led to the slight accumulation of SUMOylated αKNL2 in Arabidopsis. We further showed that SUMOylation of αKNL2 promotes its binding to CENH3 and controls protein stability. Our findings show that C-terminal SUMOylation of αKNL2 is crucial for its centromeric localization, interaction with CENH3, and kinetochore assembly, emphasizing the significance of post-translational modifications in chromosome segregation and cell division in plants.

## Introduction

Centromeres are essential chromosomal regions that ensure the accurate segregation of chromosomes during cell division. They serve as the assembly sites for the kinetochore, a multiprotein complex that interacts with spindle microtubules to facilitate chromosome movement. CENH3, also referred to as centromeric histone H3 (known as CENP-A in humans), acts as a marker for active centromeres and is crucial for kinetochore establishment (Talbert *et al*., 2002, Naish and Henderson, 2024). KINETOCHORE NULL2 (αKNL2) is a key kinetochore protein that plays a pivotal role in CENH3 loading to centromeres and kinetochore assembly. It is part of the CCAN (Constitutive Centromere-Associated Network) complex, which serves as a structural bridge between the CENH3 nucleosomes and outer kinetochore components. Arabidopsis αKNL2 contains a conserved SANTA domain in its N-terminal part, similar to KNL2 in other organisms (Zhang *et al*., 2006). Furthermore, most vertebrate and plant KNL2 homologs have been found to possess a CENPC-k motif at their C-terminus, which facilitates binding to CENH3 nucleosomes (French *et al*., 2017, Hori *et al*., 2017, Sandmann *et al*., 2017). The deletion of the CENPC-k motif or the mutation of its single conserved amino acid abolishes the centromeric localization of αKNL2 (Sandmann *et al*., 2017). The reduced levels of CENH3 protein in a *knl2* knockout mutant have been linked to micronuclei in pollen tetrads, anaphase bridges during mitosis, and 30% seed abortion (Lermontova *et al*., 2013). In *Arabidopsis thaliana*, full-length αKNL2 cannot be stably overexpressed due to its targeted degradation via the ubiquitin-proteasome system, and this process is orchestrated by APC/C^CDC20^-mediated ubiquitination of αKNL2, which is essential for maintaining kinetochore function and centromere integrity in *Arabidopsis* (Lermontova *et al*., 2013, Kalidass *et al*., 2025). This highlights the importance of post-translational modification of αKNL2 for mitotic fidelity and plant development.

Protein post-translational modifications (PTMs) play key roles in virtually all cellular processes. PTMs include small chemical modifications like phosphorylation, ubiquitination and SUMOylation. Ubiquitin polymers are well known for their classical role in targeting proteins to the proteasome for degradation. Although a classic fate of ubiquitinated proteins is proteasome-mediated degradation, the Small Ubiquitin-like Modifier (SUMO) is often associated with modulating protein-protein interactions via its non-covalent interaction with a SUMO-interacting motif (SIM), which can also influence protein localization (Mahajan *et al*., 1998, Matunis *et al*., 1998, Pichler *et al*., 2005). In addition, SUMO shares structural similarity with ubiquitin and, during the process of SUMOylation, is covalently attached to lysine residues in substrate proteins through a cascade of E1 activating, E2 conjugating, and E3 ligase enzymes. In Arabidopsis, there are four SUMO isoforms: SUMO1, SUMO2, SUMO3, and SUMO5. Among these, SUMO1 and SUMO2 are nearly identical and function redundantly, while SUMO3 and SUMO5 exhibit more specialized roles. Under normal physiological conditions, SUMOs primarily function as monomers, mediating signaling processes. Their activity is tightly regulated by SUMO-specific proteases, which remove SUMO modifications from target proteins in a process known as deSUMOylation. SUMO deconjugation is performed by the conserved ULP/SENP (ubiquitin-like protease/sentrin-specific protease) family of SUMO-specific proteases. Arabidopsis ULPs comprise a family of at least seven elements (ESD4, ULP1a/ELS1, ULP1b, ULP1c/OTS2, ULP1d/OTS1, ULP2a, and ULP2b), which may result in both specificity and redundancy within the SUMO pathway (Chosed *et al*., 2006, Colby *et al*., 2006). ESD4 and ULP1a were previously associated with the control of flowering time and plant development (Murtas *et al*., 2003, Hermkes *et al*., 2011). ULP1c and ULP1d have been implicated in salt stress responses, in the modulation of SA signaling, and the deSUMOylation of phytochrome-B (Conti *et al*., 2008, Conti *et al*., 2014, Sadanandom *et al*., 2015, Bailey *et al*., 2016).

SUMOylation is a crucial modification of chromatin proteins, including histone H3. Centromere- and kinetochore-associated factors are reported to be enriched amongst SUMOylated proteins in several species (Azuma *et al*., 2003, Montpetit *et al*., 2006, Zhang *et al*., 2008, Mukhopadhyay *et al*., 2010, Li *et al*., 2016), and both SUMO E3 ligase and SUMO protease enzymes have been found to colocalize with kinetochores (Ban *et al*., 2011, Cubeñas-Potts *et al*., 2013, Suhandynata *et al*., 2019). Conversely, recent evidence suggests that SUMOylation of kinetochore protein Nuf2 is required to promote the recruitment of the SIM- containing centromere-associated protein CENP-E, which is essential for the proper alignment of chromosomes in metaphase (Subramonian *et al*., 2021). In *Arabidopsis thaliana*, conserved AAA-ATPase molecular chaperone, CDC48/p97 interacts with SUMOylated CENH3 to remove it from centromeres, leading to the disruption of centromeric heterochromatin and the activation of rRNA genes (Mérai *et al*., 2014). Nevertheless, previous reports show that the SUMO protease (SENP6) mediates MIS18BP1/HsKNL2 deSUMOylation to regulate CENP-A loading in humans. SENP6 depletion leads to RNF4-mediated degradation of HsKNL2 and, consequently, the failure of CENP-A to accumulate at the centromere (Fu *et al*., 2019, Liebelt *et al*., 2019). HsKNL2 was co-modified by SUMO and ubiquitin, which was subsequently identified as a novel target of SUMO-targeted ubiquitin ligase (RNF4). These findings provide evidence that SUMO-ubiquitin crosstalk regulates HsKNL2 during mitosis (Cuijpers *et al*., 2017). Thus, these data indicate that the correct timing and specificity of kinetochore localization are regulated by SUMOylation. Nonetheless, the role of SUMOylation at centromeres in plants appears to be intricate and remains to be fully elucidated.

In this study, we uncovered the critical role of Arabidopsis αKNL2 SUMOylation in its localization and interaction with CENH3 at centromeres. We demonstrated the interaction of SUMO3 and ULP1d with αKNL2 using yeast two-hybrid library screening, bimolecular fluorescence complementation (BiFC), and co-immunoprecipitation (Co-IP) approaches. Additionally, we identified SUMOylation and SIM sites in the αKNL2-C, which are essential for its proper centromeric localization. Among these, SUMOylation at Lys474 and Lys511 is crucial for αKNL2 SUMOylation. The accumulation of the SUMO mutant variant of αKNL2 led to defects in growth, fertility, and mitosis. The in vitro and in vivo SUMOylation assay showed that αKNL2-C is modified by SUMO1 or SUMO3. Additionally, we found that the *ulp1d-2* mutant resulted in the accumulation of SUMOylated αKNL2. Our study also revealed that the αKNL2-C interacts with the CENH3, while the interaction was disrupted in the SUMOylation-deficient αKNL2 mutant.

## Results

### SUMO3 and ULP1d interact with the C-terminus of αKNL2

Affinity purification-mass spectrometry (AP-MS) and yeast two-hybrid (Y2H) library screening were employed to identify proteins interacting with αKNL2. For this purpose, the full-length Arabidopsis αKNL2 protein (αKNL2, 1 to 598 aa), along with its N-terminal (αKNL2-N; 1 to 363 aa) and C-terminal (αKNL2-C; 364 to 599 aa) fragments were used (Figure 1A). Pathway enrichment analysis revealed that αKNL2 is involved in processes such as post-translational modifications (PTMs), metabolism, and transcription (Kalidass *et al*., 2025). To further investigate the functional roles of these interacting proteins, gene ontology (GO) enrichment analysis was performed, focusing on biological processes (BP) and molecular functions (MF) related to PTMs. These PTMs include BP and MF categories such as ubiquitination, SUMOylation, nucleocytoplasmic transport, and RNA export from nucleus (Supplementary Figure 1). Furthermore, protein interaction network analysis indicated that αKNL2 is regulated by SUMOylation pathways, leading to the identification of a specific subnetwork of SUMOylation-related αKNL2 interactors (Figure 1B).

**Figure 1.**
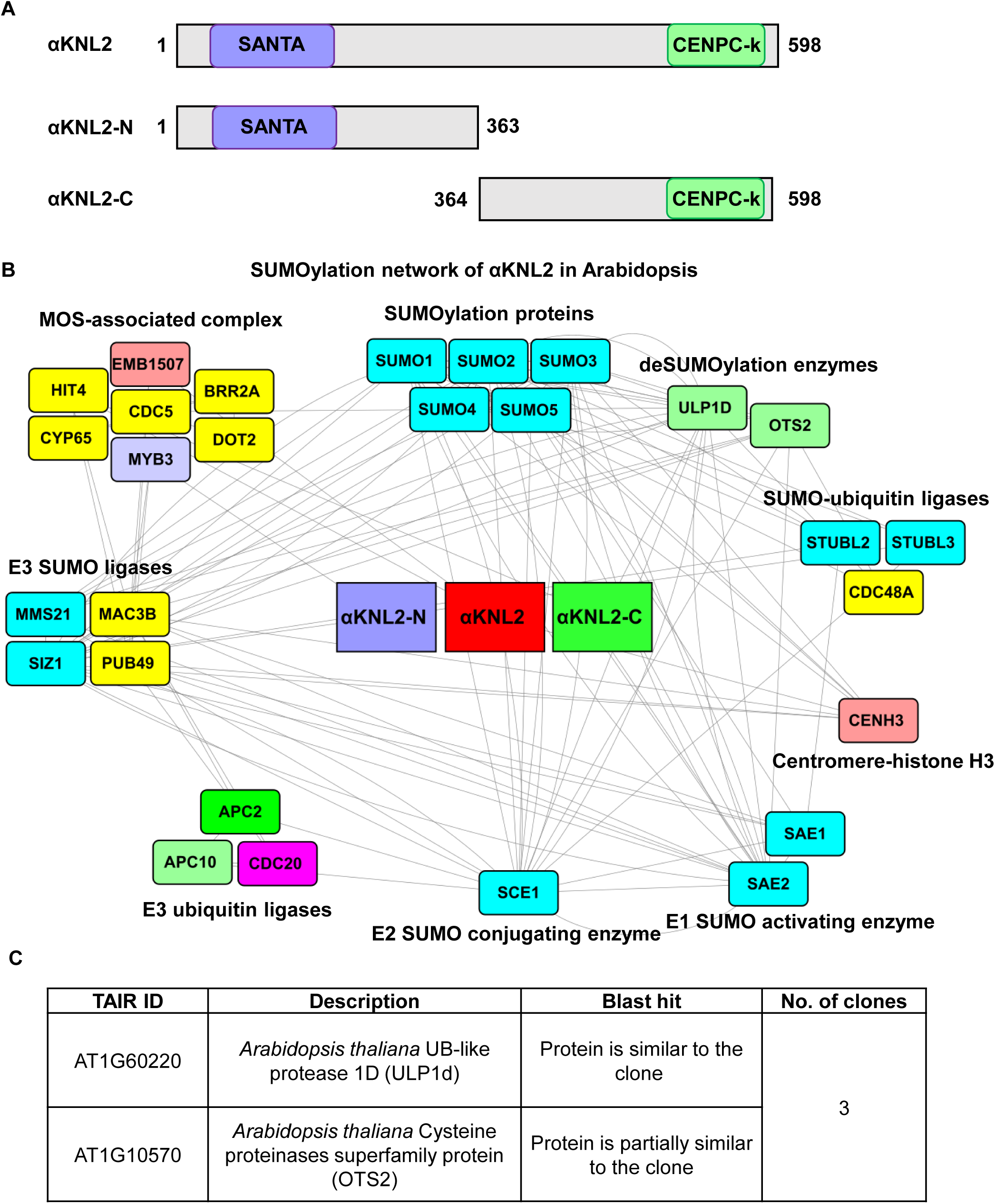
αKNL2 interactome reveals associations with the SUMOylation machinery in Arabidopsis. **(A)** The schemata illustrates the domain organization of the αKNL2 protein (1-598 aa), highlighting its N-terminal (1-363 aa) and C-terminal regions (363-598 aa). The SANTA domain, represented by a purple box, is located in the N-terminal region and the C-terminal region contains the conserved CENPC-k motif, shown in green. **(B)** The protein-protein interaction network for αKNL2 was generated from Y2H library screening and AP-MS results. Rectangular boxes represent αKNL2 interactors, grouped by their functional annotations. Interactors identified through AP-MS are shown in yellow boxes, while those identified via Y2H are displayed in colored boxes. Proteins in blue boxes, identified through STRING, were used to connect pathways but were not identified as αKNL2 interactors. The network was constructed using STRING and Cytoscape software. **(C)** The sequencing analysis of αKNL2-C clones from the Y2H screening identified ULP1d as an interactor.

To explore the role of SUMOylation in the regulation of αKNL2, ULP1d, a SUMO protease identified through Y2H library screening of αKNL2-C (Figure 1C), and the SUMO isoforms such as SUMO1, SUMO2, SUMO3, SUMO5 were selected for further validation of their interactions with αKNL2 via BiFC analysis. In the BiFC assay, full-length αKNL2, αKNL2-N and αKNL2-C were fused to the N-terminal half of Venus (VENn), while ULP1d and the SUMO isoforms were fused to the C-terminal half of Venus (VENc), and vice versa. The interaction analysis revealed that αKNL2-C interacts with ULP1d and SUMO3 within the nucleolus (Figure 2A, Supplementary Figure 2). BiFC analysis further showed that αKNL2-C specifically interacts only with the C-terminal region of ULP1d in the nucleolus (Figure 2A). Interestingly, yeast two-hybrid screening confirmed the interaction between αKNL2-C and the C-terminal region of ULP1d (ULP1d-C; 278–585 aa). Consistently, BiFC quantification showed a strong interaction between αKNL2-C and both SUMO3 and ULP1d-C, as indicated by a high number of nuclei displaying BiFC fluorescence signals and increased fluorescence intensity (Figure 2B, C). Moreover, full-length αKNL2 and αKNL2-N showed no interaction with ULP1d or SUMO3 when fused to either half of Venus. Furthermore, none of the αKNL2 fragments interacted with SUMO1, SUMO2, and SUMO5 (Supplementary Figure 2), suggesting that SUMO3 specifically binds to αKNL2.

**Figure 2.**
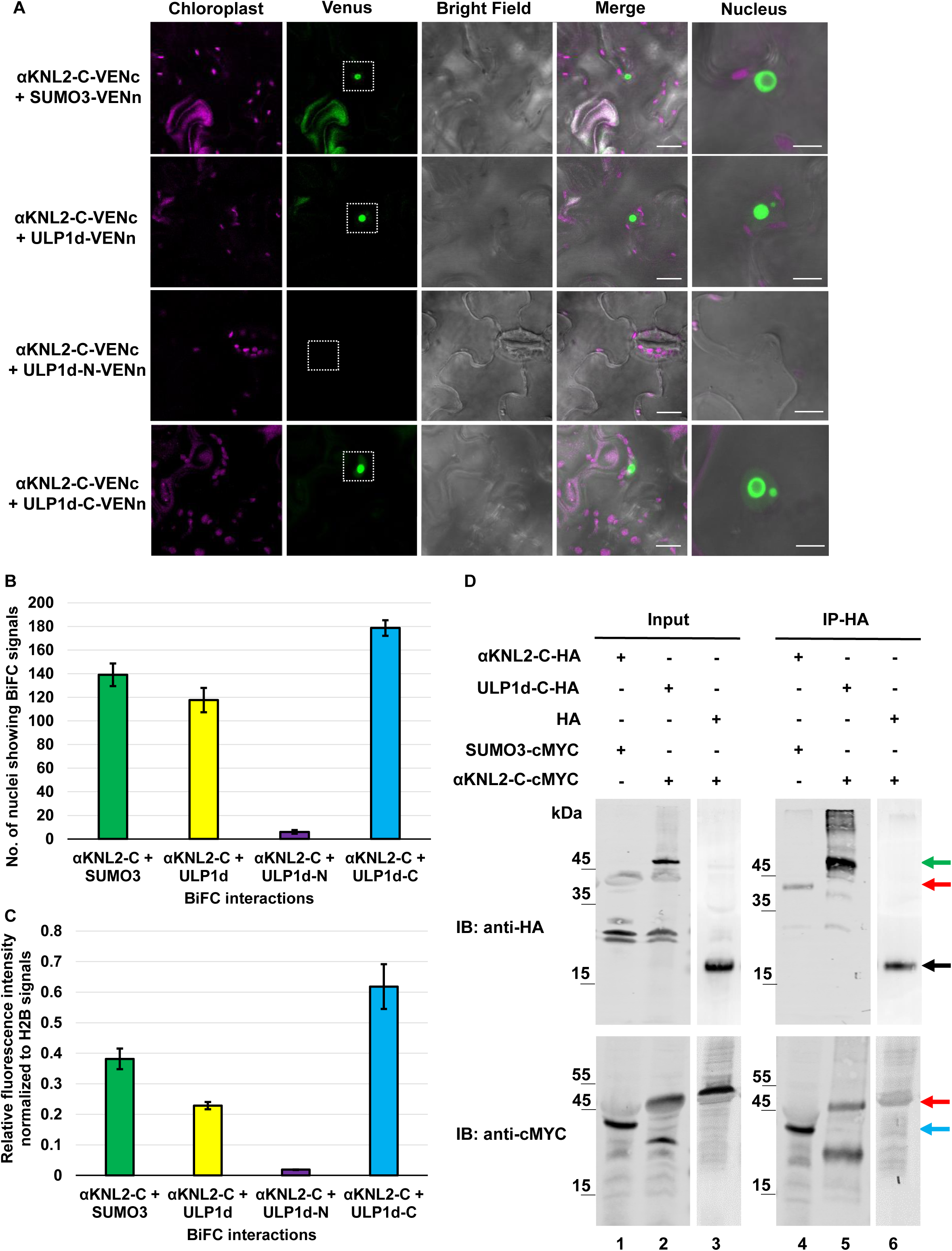
The interaction of αKNL2 C-terminus with SUMO pathway components. **(A)** BiFC analysis showing interactions between αKNL2-C fused to VENc and SUMO3, ULP1d, ULP1d-N, or ULP1d-C fused to VENn. The white dotted boxes indicate BiFC signals (Venus fluorescence). Scale bars represent 50 µm. The right panel shows a magnified view of the BiFC signals in the nucleolus. Note that the enlarged images may not always correspond to the exact same nuclei shown in the overview image. Scale bars represent 5 µm. **(B, C)** Bar graphs represent the number of nuclei showing BiFC signals (B) and the corresponding mean fluorescence intensity (C) for each interaction pair. The number of nuclei showing BiFC signals was measured in 80 mm^2^ area. The fluorescence intensity for BiFC signals were measured after normalization with H2B signals from 30 nuclei per sample (n = 30). Data are presented as mean ± SEM. **(D)** Co-IP analysis showing interactions between αKNL2-C and SUMO3 or ULP1d-C. *N. benthamiana* leaves were infiltrated with constructs encoding αKNL2-C-HA and SUMO3-cMYC (lanes 1, 4), ULP1d-C-HA and αKNL2-C-cMYC (lanes 2, 5) or HA and αKNL2-C-cMYC (lanes 3, 6). Total protein extracts were immunoprecipitated using HA magnetic beads, and samples were analyzed by immunoblotting with HA and cMYC antibodies before (Input) and after (IP) immunoprecipitation. The interactions between αKNL2-C and SUMO3 or ULP1d-C were detected by cMYC western blot, whereas no interaction was detected with the empty-HA control. The red, green, blue, and black arrows indicate the molecular weight (MW) of αKNL2-C, ULP1d-C, SUMO3, and empty-HA respectively. Abbreviations: Immunoblot (IB);Immunoprecipitation (IP).

To further validate the interaction between αKNL2 with SUMOylation pathway components, a co-immunoprecipitation (Co-IP) assay was performed. Specifically, αKNL2-C^HA^ was co-expressed with SUMO3^cMYC^, while ULP1d-C^HA^ was co-expressed with αKNL2-C^cMYC^ in *Nicotiana benthamiana* leaves. As a negative control, αKNL2-C-cMYC was co-expressed with an empty-HA vector. In all cases, total protein extracts were subjected to immunoprecipitation using HA magnetic beads.. Subsequent western blot analysis performed with an anti-cMYC antibody detected SUMO3^cMYC^ and αKNL2-C^cMYC^when co-expressed with αKNL2-C^HA^ or ULP1d-C^HA^, but not in the empty-HA control (Figure 2D). These findings corroborate the results of BiFC and yeast two-hybrid assays, confirming the interactions between αKNL2-C and SUMO3 or ULP1d.

### SUMOylation sites in the C-terminus of αKNL2 regulate its centromere targeting

The increasing identification of SUMO sites in eukaryotic cells has enabled the development of computational tools, such as GPS-SUMO (http://sumosp.biocuckoo.org/), to predict potential SUMOylation targets. Using this tool, three lysine residues K378, K474, and K511, were identified as potential SUMOylation sites in the αKNL2-C of *Arabidopsis*. Additionally, two SUMO interaction motifs (SIMs), located at residues 547-551 and 568-572 within the C-terminal region, were identified. Sequence alignment of αKNL2 homologs from Brassicales genomes confirmed the conservation of these SUMOylation and SIM sites (Figure 3A, Supplementary Figure 3A-C).

**Figure 3.**
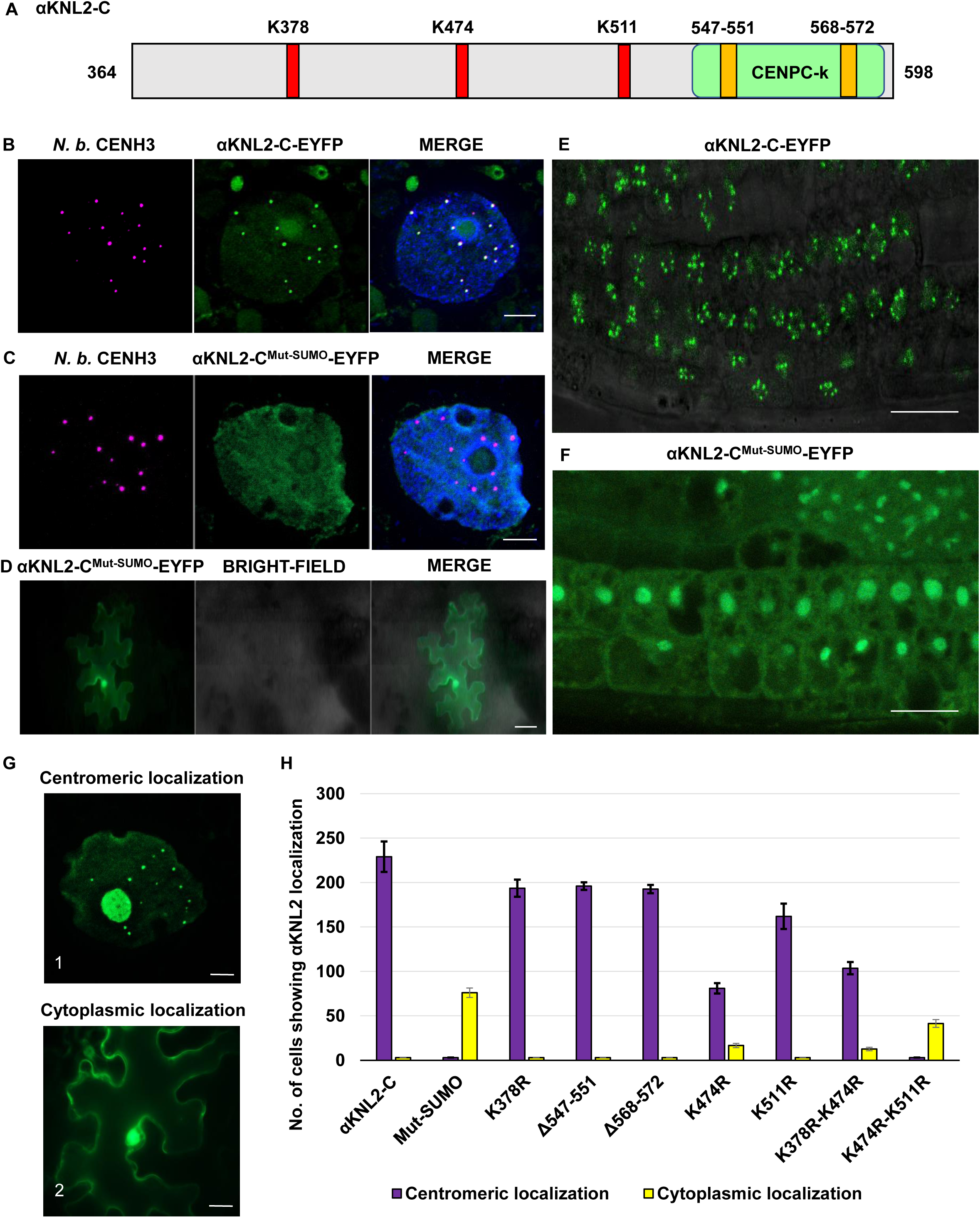
SUMOylation-deficient mutant of αKNL2-C disrupts its centromere targeting. **(A)** The C-terminal part of αKNL2 (αKNL2-C; 364-598 amino acids) contains three conserved lysine (K) residues at positions K378, K474, and K511 (red boxes), as well as two SUMO interaction sites (orange boxes; 547–551 and 568–572). **(B)** Co-localization of αKNL2-C-EYFP (green) with CENH3 (magenta) in *N. benthamiana*, indicating centromere-specific signals. Scale bars represent 5 µm. **(C, D)** The localization patterns of the SUMOylation-deficient mutant of αKNL2 (αKNL2-C^Mut-SUMO^-EYFP) in *N. benthamiana* leaves. The construct showed nucleoplasmic signals (C) that did not co-localize with *N. benthamiana* CENH3 at centromeres and cytoplasmic localization (D). Scale bars represent 5 µm. (C), 50 µm (D). **(E, F)** The localization of αKNL2-C-EYFP (E) and αKNL2-C^Mut-SUMO^-EYFP (F) in *Arabidopsis* root tips, resembling the patterns observed in *N. benthamiana*. Scale bars represent 10 µm. **(G)** The centromere-specific signals (panel 1) and cytoplasmic localization patterns (panel 2) were observed for individual SUMOylation and SIM site mutations in αKNL2-C. Scale bars represent 5 µm. **(H)** Quantitative analysis of cells displaying distinct fluorescence patterns from (G), comparing SUMO mutant variants with the αKNL2-C as a control. The number of cells displaying αKNL2 localization was quantified within an 80 mm^2^ area. For some constructs, zero values were plotted as 3 to aid visualization. Data are presented as mean ± standard error of the mean (SEM).

Consequently, a SUMOylation-deficient mutant of αKNL2-C was generated, in which these three lysine residues were substituted with arginine (K→R), and the SIMs were deleted (αKNL2-C^Mut-SUMO^). The mutated protein variant was fused to EYFP and expressed in *N. benthamiana* plants under the 35S promoter. The co-localization assays in *N. benthamiana* showed that wild-type αKNL2-C co-localized with CENH3 at centromeres, whereas the SUMOylation-deficient mutant of αKNL2 failed to localize to centromeres and instead accumulated in the nucleoplasm and cytoplasm (Figure 3B–D). To further examine the in vivo localization of αKNL2-C^Mut-SUMO^ in *A. thaliana*, stable transgenic lines expressing either αKNL2-C-EYFP and αKNL2-C^Mut-SUMO^-EYFP fusion constructs were generated. Root tip analysis of at least three independent T2 lines expressing αKNL2-C^Mut-SUMO^-EYFP showed a clear loss of αKNL2-specific centromeric signals compared to the unmutated variant. Additionally, the fluorescence was largely distributed throughout the cytoplasm and nucleoplasm, and in some cases, the nucleolus. (Figure 3E, F). Western blot analysis using an anti-GFP antibody confirmed comparable αKNL2 protein levels across all three independent lines for both the wild-type and SUMOylation-deficient constructs, indicating that the observed localization differences are not due to differences in protein expression (Supplementary Figure 4). Additionally, BiFC analysis of the SUMO-mutant of αKNL2-C showed no fluorescence when co-expressed with ULP1d, ULP1d-C or SUMO3 constructs, indicating that the SUMOylation sites and SIMs are crucial for αKNL2-C binding to SUMO3/ULP1d partners (Supplementary Figure 5A-D). These findings demonstrate that SUMOylation and/or SUMO interaction plays a crucial role in the centromeric localization of αKNL2.

To further dissect the roles of SUMOylation and SUMO interaction, site-directed mutagenesis was performed on each of the three predicted SUMOylation sites and two SIMs in αKNL2-C. The resulting mutated constructs were fused to EYFP and transiently expressed in *N. benthamiana* leaves. Surprisingly, all single lysine mutated constructs displayed centromere-specific localization. Specifically, αKNL2-C^K378R^-EYFP, αKNL2-C^K511R^-EYFP, αKNL2-C^Δ547-551^-EYFP, and αKNL2-C^Δ568-572^-EYFP retained centromeric localization, whereas αKNL2-C^K474R^-EYFP showed both centromeric and cytoplasmic localization (Figure 3G). Therefore, αKNL2 double mutants were generated targeting lysine residues 378 and 474 (K378R/K474R) or 474 and 511 (K474R/K511R). The localization pattern of αKNL2-C^K378R/K474R^-EYFP resembled that of the K474R single mutant. In contrast, αKNL2-C^K474R/K4511R^-EYFP was predominantly mislocalized to the cytoplasm, with fewer cells showing centromere-associated signals (Figure 3G). Quantification of the fluorescence patterns revealed that αKNL2-C^K378R^-EYFP, αKNL2-C^K511R^-EYFP, αKNL2-C^Δ547-551^-EYFP, and αKNL2-C^Δ568-572^-EYFP displayed centromeric localization (198–211 nuclei per 80 mm²) comparable to the wild-type αKNL2-C-EYFP (220–232 nuclei per 80 mm²). In contrast, αKNL2-C^K474R^-EYFP, and αKNL2-C^K378R/K474R^-EYFP exhibited partial loss of centromeric localization (81–103 nuclei per 80 mm²) along with cytoplasmic signals (12–16 cells per 80 mm²). Notably, αKNL2-C^K474R/K511R^-EYFP showed nucleoplasmic and cytoplasmic signals (42 cells per 80 mm²), similar to the αKNL2-C^Mut-SUMO^-EYFP pattern (76 cytoplasmic cells per 80 mm²) (Figure 3H). In addition, BiFC analysis of αKNL2-C^K474R/K511R^ showed no detectable fluorescence when co-expressed with ULP1d, ULP1d-C, or SUMO3, unlike other single or double lysine mutants (Supplementary Figure 5E). These findings suggest that SUMOylation at Lys474 and Lys511 is critical for both interaction with SUMO pathway components and centromeric targeting of αKNL2-C.

### The SUMOylation-deficient mutant of αKNL2 impairs plant development and mitosis

Given that the SUMOylation-deficient mutant of αKNL2 showed disrupted centromere targeting in *N. benthamiana* and Arabidopsis, its effects on the plant growth and development were investigated. Transgenic Arabidopsis plants expressing αKNL2-C-EYFP did not exhibit any phenotypic differences compared to wild-type (Col-0) plants (Lermontova *et al*., 2013). Therefore, the αKNL2-C-EYFP line was used as a control to assess the impact of the SUMOylation-deficient αKNL2 mutant. Following fluorescence screening of twelve independent lines, three transgenic lines exhibiting reproducible and uniform expression were selected for further study. Analysis of three independent transgenic lines expressing the αKNL2-C^Mut-SUMO^-EYFP fusion construct revealed up to an average of 28.14 % reduction in root length compared to αKNL2-C-EYFP plants (Figure 4A, B). Additionally, plants expressing αKNL2-C^Mut-SUMO^ exhibited significant differences in vegetative growth and development (Figure 4C). Previous studies have demonstrated that the *αknl2* knockout mutant, as well as lines expressing degradation-resistant variants of αKNL2 with mutations in ubiquitination sites, display mitotic defects and reduced fertility (Lermontova *et al*., 2013, Kalidass *et al*., 2025). Based on these findings, we hypothesized that overexpression of αKNL2-C^Mut-SUMO^-EYFP, which fails to localize to centromeres in Arabidopsis may lead to mitotic abnormalities.

**Figure 4.**
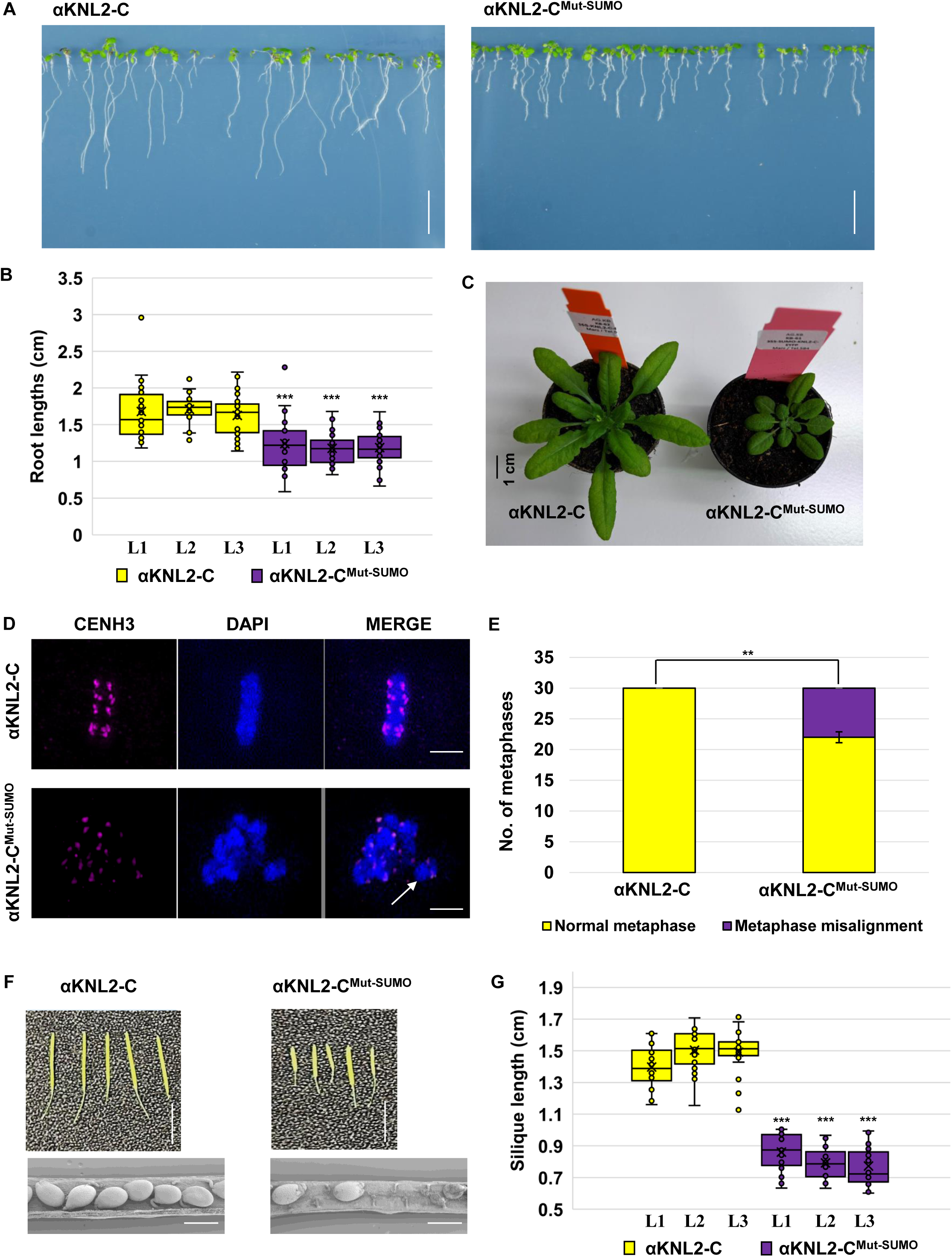
Phenotypic analysis of the SUMOylation-deficient mutant of αKNL2. **(A)** Root growth phenotype of 7-day-old *Arabidopsis* seedlings expressing αKNL2-C^Mut-SUMO^-EYFP compared to αKNL2-C-EYFP. Scale bars represent 1 cm. **(B)** Box plot showing primary root length in αKNL2-C-EYFP and αKNL2-C^Mut-SUMO^-EYFP seedlings. Seven-day-old seedlings from three independent transgenic lines per construct were analyzed (n = 25 seedlings per line). Box plots display the median (horizontal line), interquartile range (box), and data range (whiskers), with individual data points overlaid. The mean is marked by an "×". αKNL2-C^Mut-SUMO^-EYFP lines exhibited shorter primary roots compared to αKNL2-C-EYFP lines. Statistical significance was determined using Welch’s t-test; *** indicates p < 0.005. **(C)** Phenotypic comparison of 5-week-old plants expressing αKNL2-C^Mut-SUMO^-EYFP and αKNL2-C-EYFP grown in soil. **(D)** Mitotic metaphase images of αKNL2-C-EYFP and αKNL2-C^Mut-SUMO^-EYFP plants visualized by 3D-SIM, highlighting misaligned chromosomes (white arrows). Scale bars represent 5 µm. **(E)** Quantification of abnormal metaphases in SUMOylation-deficient mutant of αKNL2. Analysis of 30 metaphase cells per line revealed that 26% of metaphases in αKNL2-C^Mut-SUMO^-EYFP displayed misalignment. Data were represented from three independent lines and are shown as means ± SEM (n = 3). Significant differences between groups were assessed using Welch’s t-test and are indicated by *** (p < 0.05) **(F)** Comparison of silique size between αKNL2-C-EYFP and αKNL2-C^Mut-SUMO^-EYFP plants (upper panel). Scale bars represent 1 cm. Scanning electron microscopy images of siliques (lower panel). Scale bars represent 20 µm. **(G)** Box plot showing silique length in αKNL2-C-EYFP and αKNL2-C^Mut-SUMO^-EYFP plants. Siliques from three independent transgenic lines per construct were analyzed (n = 25 siliques per line). The box plots display the median (horizontal line), interquartile range (box), and data range (whiskers), with individual data points. The mean is indicated by an "×". Statistical significance was assessed using Welch’s t-test; *** denotes p < 0.005.

Consistent with this hypothesis, mitotic analysis of root tip meristems from three independent transgenic lines expressing αKNL2-C^Mut-SUMO^-EYFP revealed mitotic abnormalities. On average, 26% of the analyzed cells (8 out of 30), displayed misaligned metaphase chromosomes (Figure 4D, E, Supplementary Figure 6A). Fertility assessments further revealed impaired reproductive development in αKNL2-C^Mut-SUMO^-EYFP, as evidenced by reduced silique size compared to αKNL2-C-EYFP plants across three independent lines (Figure 4F, G). However, pollen viability was unaffected, as confirmed by Alexander staining (Supplementary Figure 6B). Furthermore, seed analysis from 10 siliques of a representative αKNL2-C^Mut-SUMO^-EYFP line showed that, on average, 20% of seeds were aborted, and 18% were shriveled. (Supplementary Figure 6B, C). These findings suggest that SUMOylation of αKNL2 is crucial for mitotic progression, and fertility in Arabidopsis.

### In vivo and in vitro SUMOylation reveals isoform-specific modification of αKNL2

To investigate whether αKNL2 undergoes SUMOylation *in planta*, total proteins were extracted from leaves of *N. benthamiana* infiltrated with constructs expressing αKNL2-C-EYFP, αKNL2-C^Mut-SUMO^-EYFP or EYFP alone. The proteins were immunoprecipitated using GFP affinity beads and analyzed by immunoblotting with either a mouse anti-GFP monoclonal antibody or a rabbit anti-SUMO3 antibody. Immunoblotting with anti-GFP detected bands corresponding to the molecular weight (MW) of the αKNL2-C-EYFP fusion protein (∼55 kDa), along with additional bands of higher MWs, suggesting post-translational modifications of αKNL2-C. In contrast, the αKNL2-C^Mut-SUMO^-EYFP sample exhibited reduced or absent bands at these MWs, indicating that the mutated lysines in this construct may serve as potential SUMOylation sites (Figure 5A). To examine the SUMOylation of αKNL2, specific antibodies against SUMO3 and SUMO1 were used. Immunoblotting analysis revealed that SUMO3 and SUMO1 covalently bind to αKNL2-C, forming distinct bands corresponding to higher and lower MWs. The higher MW bands (above 55 kDa) likely represent αKNL2-C modified with one or more covalently attached SUMO molecules. In contrast, the lower MW bands may result from degradation products. Interestingly, the SUMOylation-deficient mutant exhibited reduced SUMO3 binding of αKNL2-C, while SUMO1 binding was only minimally affected. However, no SUMOylation was detected in the EYFP pull-down control (Figure 5A, Supplementary Figure 7A). This confirms that the SUMO site mutations disrupted the attachment of SUMO3, with a minor effect on SUMO1 binding. This validates that the modifications observed in the wild-type αKNL2-C-EYFP were indeed due to SUMOylation of the conserved lysine residues.

**Figure 5.**
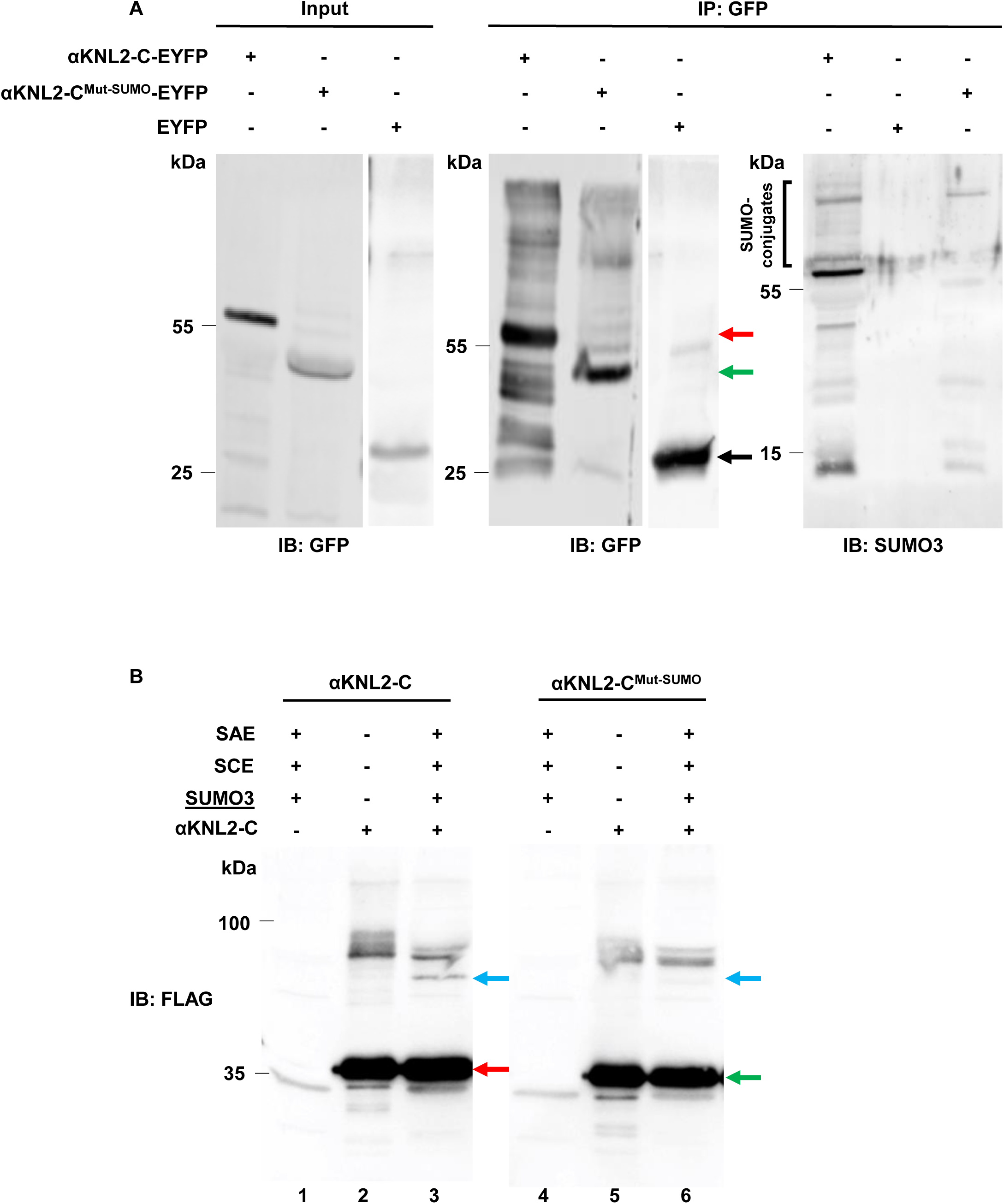
The in vivo and in vitro SUMOylation analysis of αKNL2. **(A)** The in vivo SUMOylation analysis of αKNL2 in *N. benthamiana* leaves expressing αKNL2-C-EYFP, or αKNL2-C^Mut-SUMO^-EYFP. Leaves expressing EYFP alone served as a control. Total protein extracts were immunoprecipitated using GFP beads. Input samples were analyzed with an anti-GFP antibody, while immunoprecipitated samples were probed with anti-GFP, or anti-SUMO3. Tubulin served as the loading control. The red, green, and black arrows denote the MW of αKNL2-C, αKNL2-C^Mut-SUMO^, and EYFP, respectively. SUMO conjugates were highlighted by black brackets. Abbreviations: Immunoblot (IB);Immunoprecipitation (IP). **(B)** The in vitro SUMOylation assay reactions included enzymes only (lanes 1 and 4), substrate only (lanes 2 and 5), and a mixture of enzymes and substrates (lanes 3 and 6). After incubation, samples were analyzed using SDS-PAGE, followed by immunoblotting and anti-FLAG antibody detection of αKNL2. The full, uncropped blot, including the Ponceau S loading control, is provided in Supplementary Figure 7. The red arrow represents the unmodified αKNL2-C, the green arrow marks the unmodified αKNL2-C^Mut-SUMO^ mutant, and the blue arrows indicate SUMOylated forms. Apparently, mutations of the conserved Lys residues reduced SUMOylation efficiency compared to the wild-type variant.

To complement the in vivo observations, the in vitro SUMOylation assay (Tomanov *et al*., 2022) was performed to directly evaluate the SUMOylation efficiency of the αKNL2-C and its SUMO mutant variant. Purified αKNL2-C proteins (αKNL2-C and αKNL2-C^Mut-SUMO^; Figure 5B, lanes 2 and 5, Supplementary Figure 7B) were incubated at 30°C for 2 hours with a minimal enzymatic system comprising the E1 SUMO-activating enzyme (SAE), the E2 SUMO-conjugating enzyme (SCE), and the SUMO3 isoform (Figure 5B, lanes 1 and 4). Samples were analyzed via SDS-PAGE followed by western blotting using an anti-FLAG antibody to detect both unmodified and SUMOylated forms of αKNL2. In reactions containing αKNL2-C, a distinct higher molecular weight band corresponding to the SUMOylated form was observed (Figure 5B, lane 3). In contrast, reactions with αKNL2-C^Mut-SUMO^ exhibited significantly reduced SUMOylation levels, as indicated by the much weaker signal of the SUMOylated band (Figure 5B, lane 6). The result of the in vitro assay aligns with in vivo findings, further substantiating that SUMOylation by SUMO3 is a key post-translational modification regulating αKNL2 function. Importantly, the inability of αKNL2-C^Mut-SUMO^ to be SUMOylated by SUMO3 highlights the critical role of conserved SUMOylation sites, particularly K474 and K511 residues. To further investigate the role of different SUMO isoforms, similar *in vitro* reactions were conducted with the SUMO1 isoform. Interestingly, in this setup, no significant reduction in SUMOylation was observed for αKNL2-C^Mut-SUMO^ (Supplementary Figure 7C, left panel, lanes 3 and 7), indicating that SUMO1 can SUMOylate αKNL2^Mut-SUMO^ even in the absence of conserved SUMOylation sites. This suggests that SUMO1 may modify alternative, less conserved sites on αKNL2. In addition, SUMO1 conjugation was enhanced by E3-ligase NSE2, while SUMO3 was not (Supplementary Figure 7, compare lanes 3 and 4 or 7 and 8). These results highlight the potential for differential regulation of αKNL2 by distinct SUMO isoforms and SUMO-ligases. While SUMO3-dependent SUMOylation appears to require the conserved sites in the C-terminal region of αKNL2, SUMO1-mediated modification occurs independently of these sites, underscoring the functional diversity of SUMO isoforms in regulating αKNL2.

To validate and complement the in vitro SUMOylation results, we performed mass spectrometry (MS) analysis to identify the specific lysine residues in αKNL2-C modified by SUMO1 and SUMO3. For this purpose, wild-type αKNL2-C protein was subjected to in vitro SUMOylation reactions using either SUMO1 or SUMO3 in the presence of the SUMO-E3 ligase NSE2. As a negative control, we included the unmodified wild-type αKNL2-C protein incubated under the same conditions without the SUMOylation machinery. The MS analysis revealed that SUMO1 modified several lysine residues, including K424, K445, K511, K540, K572, K583, K585, and K598, indicating extensive SUMOylation across the C-terminal region. In contrast, SUMO3-dependent modification was detected only at K424, K511, and K598. These findings are consistent with our in vitro assay results (Supplementary Figure 7), which showed stronger SUMOylation by the SUMO1 isoform than SUMO3. Notably, K511, a lysine residue predicted in silico (via GPS-SUMO), was confirmed by MS, while another predicted site, K474, was not identified, possibly due to technical limitations or preferential modification of other residues in vitro. These findings demonstrate that αKNL2 is SUMOylated in a site- and isoform-specific manner, with SUMO3-dependent modification requiring conserved C-terminal lysines.

### SUMO conjugation on αKNL2 increases upon ULP1d knockout in Arabidopsis

ULP1d, a deSUMOylation enzyme, was identified as an interactor of αKNL2 through Y2H screening. To investigate the role of ULP1d in the deSUMOylation of αKNL2, we utilized the previously characterized T-DNA insertion mutant line *ulp1d-2* (SALK_022798) (Castro *et al*., 2016). The knockout of the ULP1d gene in homozygous *ulp1d-2* mutants was confirmed via RT-PCR analysis (Supplementary Figure 8A). Homozygous *ulp1d-2* plants exhibited distinct vegetative development phenotypes compared to heterozygous mutants and wild-type plants, consistent with previous reports (Supplementary Figure 8B).

We hypothesized that if ULP1d mediates the deSUMOylation of αKNL2, the absence of ULP1d would result in increased SUMOylation levels of αKNL2. To test this, we introduced the fusion constructs αKNL2-C-EYFP and αKNL2-C^Mut-SUMO^-EYFP into the *ulp1d-2* mutant background. Three independent T2 transgenic Arabidopsis lines expressing either αKNL2-C-EYFP or αKNL2-C^Mut-SUMO^-EYFP in the wild-type (Col-0) and *ulp1d-2* background were analyzed. Examination of root tips revealed that centromeric localization of αKNL2-C was abolished in *ulp1d-2* mutants and was primarily confined to the nucleolus, contrasting with its localization in wild-type plants (Figure 6A, Supplementary Figure 8C). This finding underscores the role of SUMOylation in the centromeric targeting of αKNL2. Conversely, αKNL2-C^Mut-SUMO^-EYFP localized predominantly to the nucleoplasm and cytoplasm, and occasionally to the nucleolus, mirroring its behaviour in the wild-type background (Figure 6A, Supplementary Figure 8C). Transgenic plants expressing the αKNL2-C^Mut-SUMO^-EYFP fusion construct exhibited vegetative growth defects, particularly in shoot development, in comparison to *ulp1d-2* mutants expressing the αKNL2-C-EYFP construct, as well as *ulp1d-2* mutants, and wild-type plants (Supplementary Figure 8D). The RT-qPCR analysis confirmed that similar *αKNL2* transcript levels across all three independent lines for both the wild-type and SUMOylation-deficient constructs in wild-type and *ulp1d-2* backgrounds, indicating that the observed phenotypic differences are not due to altered expression of *αKNL2* (Supplementary Figure 9A).

**Figure 6.**
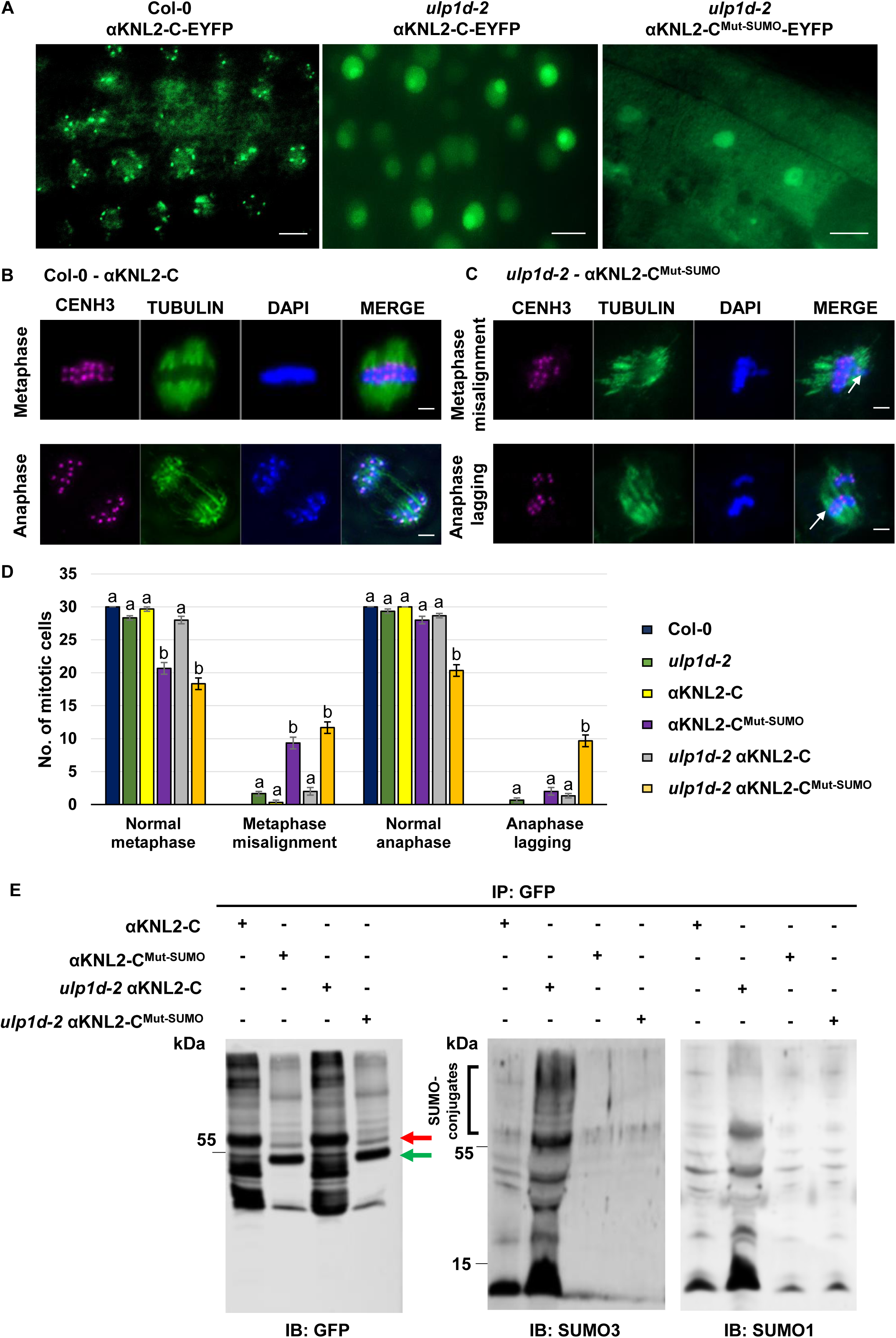
The accumulation of αKNL2 at centromeres is impaired in the absence of ULP1d. **(A)** The Arabidopsis root tips showing the localization of αKNL2-C-EYFP in wild-type, *ulp1d-2*, and αKNL2-C^Mut-SUMO^-EYFP in *ulp1d-2* mutant plants. Scale bars represent 10 µm. **(B, C)** Mitotic metaphases and anaphases of αKNL2-C-EYFP and αKNL2-C^Mut-SUMO^-EYFP in wild-type and *ulp1d-2* plants, showing normal **(B)** or misaligned and mis-segregated chromosomes shown in white arrows (n = 30) **(C)**. Scale bars represent 5 µm. Chromosomes were stained with anti-CENH3 (magenta) and anti-tubulin (green), with DAPI as a counterstain. **(D)** Abnormal metaphase and anaphase events from (B, C) were quantified. A total of 30 cells per line were analyzed, with data derived from three independent lines. Results are shown as mean ± SEM (n = 3). Significant differences are marked by lowercase letters based on ANOVA and Tukey’s multiple comparison tests (p < 0.005). **(E)** SUMOylation assay of αKNL2 in wild-type and *ulp1d-2* mutant plants. Total protein extracts were immunoprecipitated using GFP beads. Immunoprecipitated samples were analyzed with anti-GFP, anti-SUMO3 or anti-SUMO1. The red and green asterisk indicates the MW of αKNL2-C and αKNL2-C^Mut-SUMO^, respectively. SUMO conjugates were highlighted by black brackets, respectively. Abbreviations: IB, Immunoblot; IP, Immunoprecipitation.

Analysis of root tip meristems of these plants revealed significant mitotic abnormalities, including misaligned metaphases and mis-segregated chromosomes during anaphase (Figure 6B). The average frequency of mitotic abnormalities was 30-36% in αKNL2-C^Mut-SUMO^-EYFP and 4-6% in αKNL2-C-EYFP within the *ulp1d-2* background. Similarly, 6-30% of abnormalities were observed in αKNL2-C^Mut-SUMO^-EYFP compared to 0-1% in αKNL2-C-EYFP within the wild-type background. In contrast, minimal or no mitotic abnormalities were observed in *ulp1d-2* (5%) and wild-type plants (0%) (Figure 6C). To quantify αKNL2 SUMOylation levels in the *ulp1d-2* mutant relative to the wild-type, total proteins were extracted from plants expressing either αKNL2-C-EYFP or αKNL2-C^Mut-SUMO^-EYFP. Western blot of input samples with anti-SUMO3 showed increased SUMOylation in the *ulp1d-2* mutants compared to wild-type plants (Supplementary Figure 9B). Further, immunoprecipitation with GFP beads followed by immunoblotting with anti-GFP, anti-SUMO3, and anti-SUMO1 antibodies confirmed the slight accumulation of SUMOylated αKNL2-C in *ulp1d-2*. Interestingly, immunoprecipitated αKNL2-C^Mut-SUMO^-EYFP samples displayed similar patterns within wild-type and *ulp1d-2* backgrounds (Figure 6D). These findings demonstrate ULP1d as a critical protease required for the deSUMOylation of αKNL2, enabling its proper centromeric localization and functional roles in plant development.

### αKNL2 SUMOylation is required for its association with CENH3

*Arabidopsis* αKNL2 functions as a licensing factor for loading CENH3 at centromeres and is essential for kinetochore assembly. However, direct interaction between αKNL2 and CENH3 in *Arabidopsis* had not been demonstrated previously. In our AP-MS experiment, CENH3 was co-precipitated with the C-terminal region of αKNL2. To validate this association, we performed BiFC and Co-IP experiments using constructs encoding full-length αKNL2 and its N- and C-terminal regions with CENH3. The BiFC assay revealed that αKNL2-C^VENc^ specifically interacts with CENH3^VENn^ within the nucleus (Figure 7A). In contrast, neither the full-length αKNL2 nor its N-terminal region showed any detectable interaction with CENH3, even after treatment with the proteasome inhibitor MG115 (data not shown). These results demonstrate that the C-terminal region of αKNL2 interacts mainly with CENH3.

**Figure 7.**
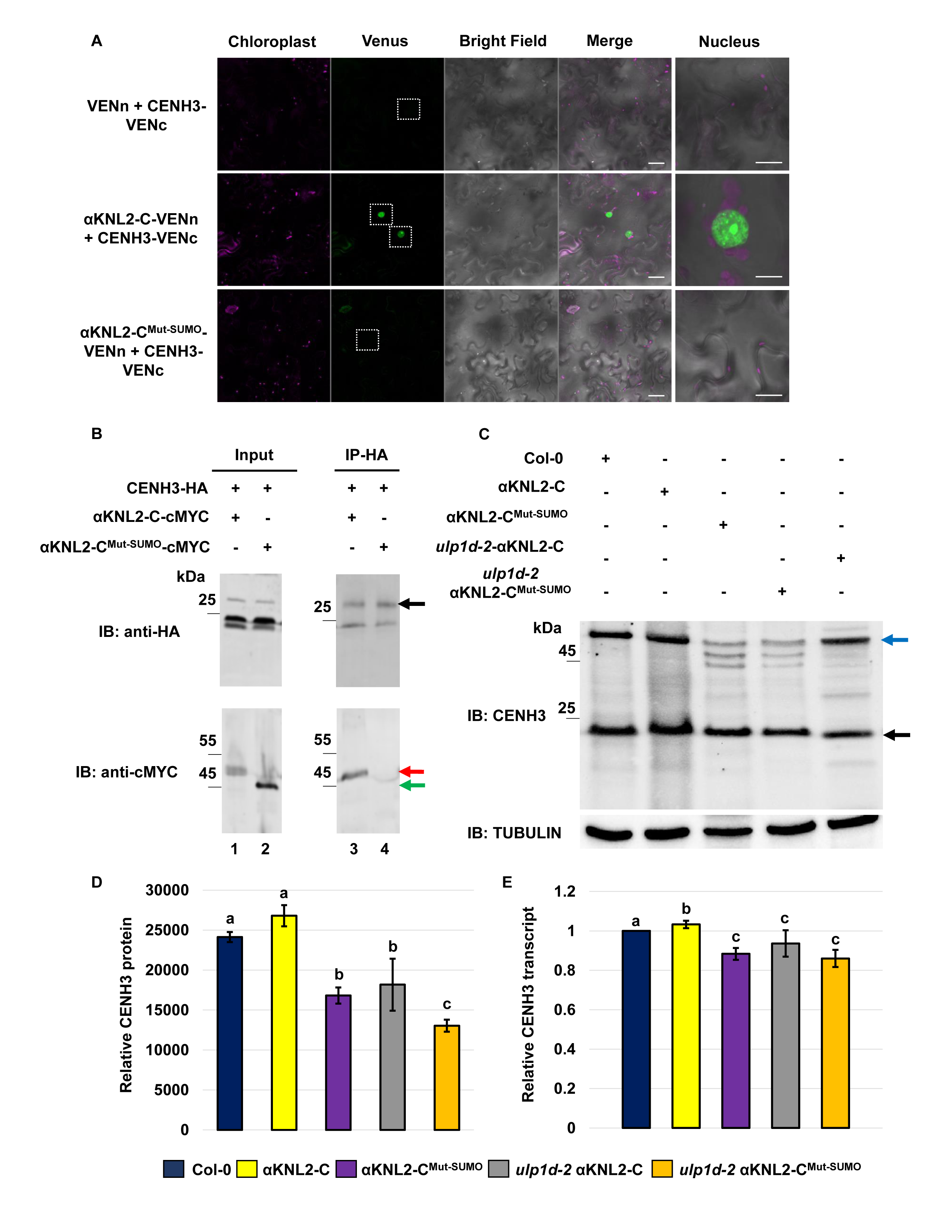
The SUMOylation of αKNL2-C is required for its interaction with the CENH3. **(A)** Confocal microscopy images of *Nicotiana benthamiana* leaf epidermal cells transiently expressing BiFC interactions of αKNL2-C fused with VENc or empty-VENc with CENH3 fused to VENn, respectively. Venus fluorescence was detected in nucleus (white dotted boxes). Scale bars represent 50 μm. The right panel displays an enlarged image of the corresponding BiFC signals in the nucleus. Note that the enlarged images may not always correspond to the exact nuclei shown in the overview image. Scale bars represent 5 µm. **(B)** Co-IP interactions between CENH3 and αKNL2-C. *N. benthamiana* leaves were infiltrated with constructs containing CENH3-HA and αKNL2-C-cMYC (lane 1, 3), CENH3-HA and αKNL2-C^Mut-SUMO^-cMYC (lane 2, 4). Total protein extracts were precipitated with HA magnetic beads and the samples were analyzed before (Input) and after (IP) immunoprecipitation by immunoblotting with HA and cMYC antibodies. The black, red, and green arrows indicate the MW of CENH3, αKNL2-C, and αKNL2-C^Mut-SUMO^, respectively. Abbreviations: IB, Immunoblot; IP, Immunoprecipitation. **(C)** Western blot analysis of Arabidopsis transgenic lines expressing αKNL2-C-EYFP or αKNL2-C^Mut-SUMO^-EYFP in Col-0 or *ulp1d-2* mutant background by anti-CENH3. CENH3 monomer and dimers are indicated by the black and blue arrows. The tubulin was used as a loading control. **(D)** Quantification of CENH3 protein levels in Arabidopsis lines shown in (C). Data are presented as mean ± SEM. Significant differences are marked by lowercase letters based on ANOVA and Tukey’s multiple comparison tests (p < 0.005). **(E)** Quantification of *CENH3* transcript levels by RT-qPCR in the same lines as in (C). Data are normalized to *ACTIN2, UBQ* expression and shown as mean ± SEM from three biological replicates. Significant differences are marked by lowercase letters based on ANOVA and Tukey’s multiple comparison tests (p < 0.005).

To investigate the factors influencing αKNL2 interaction with CENH3, we examined the role of αKNL2 SUMOylation/SIM sites using a mutant construct, αKNL2-C^Mut-SUMO^. Unlike the wild-type αKNL2-C, the SUMO mutant variant of αKNL2-C was unable to interact with CENH3 in the BiFC assay (Figure 7A, Supplementary Figure 10A), showing that SUMOylation of αKNL2-C likely facilitates its interaction with CENH3. Moreover, to confirm the interaction between αKNL2 and CENH3, a Y2H co-transfection assay was performed. The yeast strains expressing αKNL2, αKNL2-N, or αKNL2-C^BD^ as bait and CENH3^AD^ as prey did not grow on the selective TDO medium, indicating that CENH3 may not interact directly with αKNL2. CENH3^AD^ and CENH3^BD^ protein interactions were used as a positive control (Supplementary Figure 10B). This suggests that additional factors or modifications, such as SUMOylation, which are absent in yeast, might be required for the interaction. To further validate the interaction between αKNL2-C and CENH3, a Co-IP assay was performed. The fusion construct of CENH3^HA^ was co-expressed with αKNL2-C^cMYC^ or SUMOylation-deficient mutant of αKNL2-C^cMYC^. Proteins were immunoprecipitated using HA magnetic beads, and subsequent Western blot analysis with an anti-cMYC antibody detected αKNL2-C^cMYC^ co-precipitated with CENH3^HA^. Nevertheless, no interaction was observed between the SUMOylation-deficient of αKNL2-C^cMYC^ and CENH3^HA^ (Figure 7B). These findings are consistent with the BiFC results and indicate that SUMOylation and SIM sites of αKNL2-C are essential for its association with CENH3.

Consistent with these findings, western blot analysis using anti-CENH3 and anti-αKNL2 antibodies on nuclear protein extracts revealed reduced levels of endogenous CENH3 monomer (∼19 kDa) and αKNL2 (∼75 kDa) in the SUMOylation-deficient αKNL2-C mutant expressed in the wild-type background. In the *ulp1d-2* background, the αKNL2-C^Mut-SUMO^ lines exhibited an even greater reduction of CENH3 and αKNL2 proteins compared to αKNL2-C-EYFP plants. Furthermore, in wild-type plants, CENH3 is also detected as a ∼54 kDa band, likely corresponding to its incorporation into stable nucleosomal complexes containing other histones. In contrast, SUMOylation-deficient αKNL2 mutants exhibit additional bands in the ∼45–49 kDa, which may represent unstable CENH3 complexes. Notably, the transcript levels of both *αKNL2* and *CENH3* remained largely unchanged in these mutants (Figure 7C-E, Supplementary Figure 9A, 11). This suggests that αKNL2 SUMOylation regulates CENH3 and αKNL2 protein stability or deposition at centromeres rather than their transcription.

## Discussion

In Arabidopsis, αKNL2 is primarily recognized for its essential role in kinetochore assembly and CENH3 loading. As a critical regulator of mitosis, αKNL2 is tightly controlled by highly coordinated and multi-faceted regulatory mechanisms. Recent studies have begun to uncover the pathways that regulate αKNL2 activity and function in Arabidopsis (Yalagapati *et al*., 2024, Kalidass *et al*., 2025). Notably, evidence from both animal and plant systems underscores the significance of PTMs in regulating αKNL2 functionality. In this study, we demonstrated the critical role of SUMOylation in regulating αKNL2 function. Functional analyses revealed that the SUMOylation sites in αKNL2 facilitate SUMO conjugation and are indispensable for its centromeric localization and kinetochore assembly. Interestingly, expression of SUMO mutant variant of αKNL2-C in Arabidopsis wild-type background resulted in defects in chromosome alignment, growth, and fertility. Thus, this post-translational modification promotes centromere targeting of αKNL2 and interaction with CENH3, significantly enhancing its protein stability to ensure proper mitotic progression and cell division (Figure 8).

**Figure 8.**
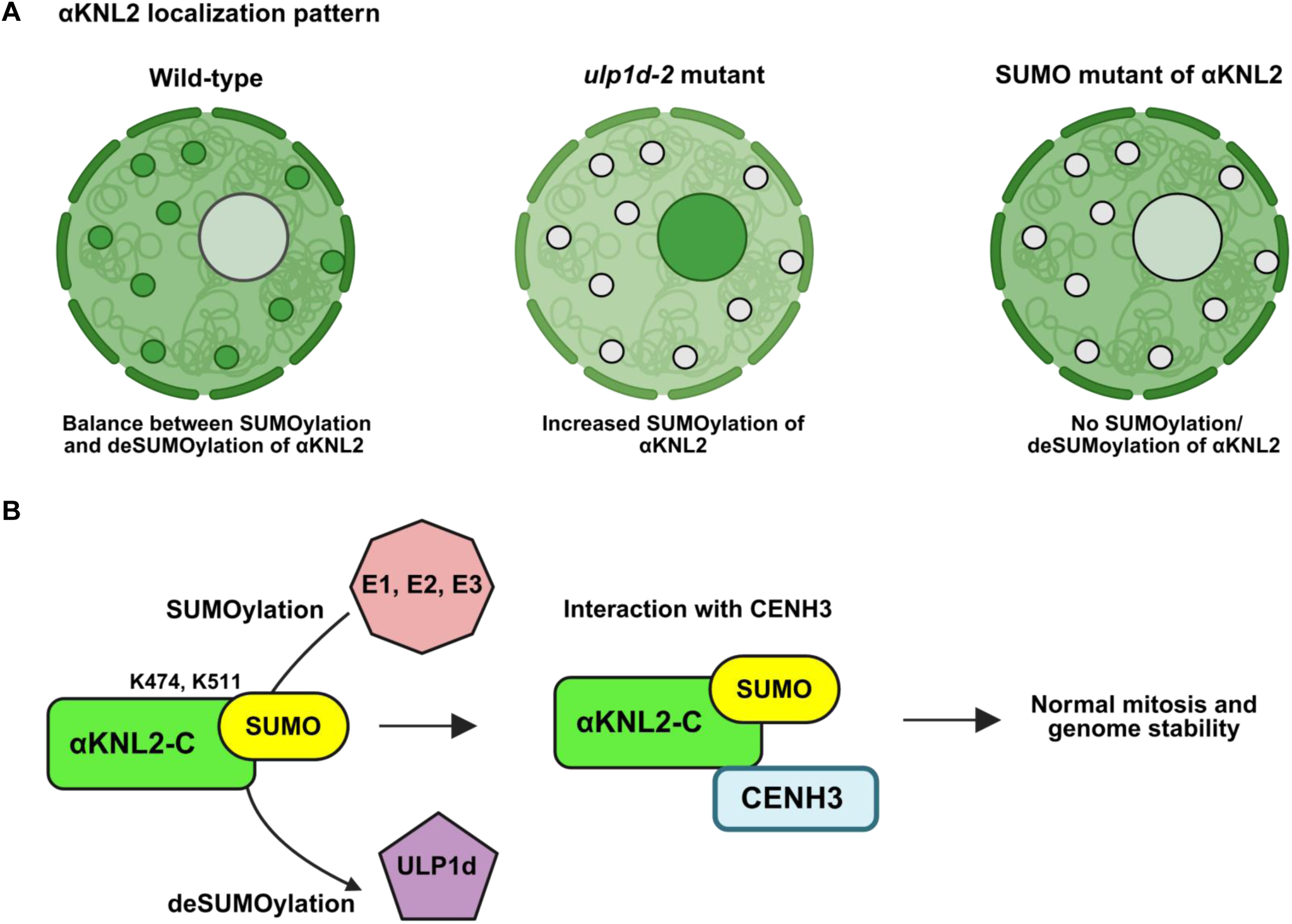
SUMOylation-dependent regulation of αKNL2 localization and its impact on centromere function. **(A)** In wild-type cells, αKNL2 (green) predominantly localizes to the centromere region and facilitates proper CENH3 deposition. In *ulp1d* mutants, αKNL2 localization is restricted to the nucleolus and is absent from the centromeres. In SUMOylation-deficient αKNL2 mutants, αKNL2-C fails to localize correctly and is entirely mislocalized to the cytoplasm. Nucleoplasmic signals were observed across all cases, with occasional weak nucleolar staining detected in both wild-type and SUMO mutant backgrounds. **(B)** Schematic representation showing the molecular mechanism of SUMOylation regulating αKNL2-C. αKNL2-C is SUMOylated at lysines K474 and K511 by the SUMO conjugation machinery (E1, E2, E3) using SUMO3/SUMO1, and this modification is reversed by the SUMO protease ULP1d. Proper SUMO cycling is essential for αKNL2-C function and efficient centromeric loading of CENH3, thereby ensuring normal mitosis and maintaining genome stability.

SUMOylation is a reversible post-translational modification that plays a pivotal role in processes such as the cell cycle, DNA repair, transcription, signal transduction, and chromatin remodeling (Müller *et al*., 2001, Hay, 2005, van den Berg and Jansen, 2023). Several key cell cycle regulators, including CENP-A (Mérai *et al*., 2014, Ohkuni *et al*., 2018, van den Berg *et al*., 2023), CENP-E (Zhang *et al*., 2008), Aurora B (Fernández-Miranda *et al*., 2010), and BUBR1 (Yang *et al*., 2012), have been identified as targets of SUMOylation. Our study provides both in vivo and in vitro evidence that αKNL2 undergoes SUMOylation, predominantly by SUMO3 and SUMO1. In mammals, SUMO1 primarily stabilizes structural proteins, whereas SUMO2 and SUMO3 are involved in the dynamic regulation of protein turnover and interactions. Similarly, in plants, SUMO1 and SUMO2 are likely to bind key centromeric proteins regulating their roles during mitosis and meiosis. SUMO3 and SUMO5 may further contribute to the modulation of centromeric protein dynamics, particularly in response to environmental stresses (van den Berg and Jansen, 2023).

Our findings indicate that αKNL2 covalently binds with both SUMO1 and SUMO3, pointing to potentially distinct regulatory roles for its SUMOylation by different SUMO isoforms. However, SUMO1 does not interact with αKNL2 in BiFC analysis, likely due to differences in the biochemical and structural properties of SUMO1 and SUMO3. The greater propensity of SUMO3 to form SUMO conjugates may enhance its binding in BiFC contexts (Chupreta *et al*., 2005, Castaño-Miquel *et al*., 2011, Park *et al*., 2011, Roy and Sadanandom, 2021). Moreover, BiFC requires a stable complex for fluorescence restoration (Miller *et al*., 2015), and SUMO3 likely forms a stronger or more stable interaction with αKNL2. In contrast, SUMO1-αKNL2 binding may be transient, structurally incompatible with BiFC detection or alternatively, it may require specific E3 ligases for its recruitment/conjugation. SUMO E3 ligases play a crucial role in facilitating SUMO conjugation and significantly enhance the efficiency of this process. An in vitro assay demonstrated that NSE2 enhances the SUMOylation of αKNL2 via SUMO1 modification. Given the canonical role of the SMC5/6 complex and its NSE2 subunit in genome stability maintenance (Aragón, 2018, Palecek, 2018), it is plausible that SUMO1 covalent binding to αKNL2 occurs through the involvement of NSE2. Increasing evidence suggests that NSE2 plays a critical role in mitosis by acting as a SUMO ligase for centromeric proteins (Andrews *et al*., 2005). Therefore, we speculate that NSE2 may function as a specific SUMO E3 ligase for αKNL2. Our future endeavors will provide a definitive answer.

Previous studies showed that αKNL2 variants fused with EYFP were found to localize in chromocenters, and occasionally in the nucleolus and nuclear bodies in *Arabidopsis* and *N. benthamiana* (Lermontova *et al*., 2013). Protein-protein interaction network analysis revealed that αKNL2 may have multiple roles at the nuclear periphery, including chromatin organization, CENH3 loading, RNA and DNA interactions, nucleotide excision repair, and regulation of nuclear transport. While association of αKNL2 with multiple partner proteins suggests roles in distinct subcellular processes, the mechanisms that govern the specificity and regulation of these interactions remain to be elucidated. We identified SUMO3 and ULP1d as specific interactors of the αKNL2-C using AP-MS, Y2H library screening, and protein interaction assays, suggesting a regulatory mechanism involving SUMOylation and deSUMOylation. This interplay between SUMO3 and ULP1d highlights a dynamic balance of SUMOylation essential for proper αKNL2 function. SUMOylation influences the subcellular localization of proteins by modulating nuclear import/export signals or promoting interactions with nucleolar targeting sequences (Müller *et al*., 2001, Wilson and Rangasamy, 2001). Proteins modified by SUMO often relocalize to nuclear or nucleolar compartments as part of their regulatory roles. Consistent with these findings, the interaction of ULP1d and SUMO3 with αKNL2 has been detected in the nucleolus and nuclear bodies, aligning with the occasionally observed localization pattern of αKNL2. Many kinetochore proteins frequently shuttle through the nucleolus for post-translational modifications, such as SUMOylation and deSUMOylation, which affect their stability, localization, and interactions. In humans, SENP3 and SENP5 are nucleolus-localized enzymes that facilitate the modification of several SUMO-2/3 substrates (Gong and Yeh, 2006).

The predictive analysis identified conserved SUMOylation sites and SIM motifs in the αKNL2-C, and mutating these conserved residues abolished centromeric localization of αKNL2-C, as evidenced by cytoplasmic mislocalization of the αKNL2-C^Mut-SUMO^ construct. However, single amino acid substitutions did not completely abrogate centromeric localization, suggesting the necessity for the presence of one or more conserved SUMO sites in αKNL2-C. Among these sites, mutation of K474, and K511 exhibited the most pronounced effect, indicating a potentially critical role for these residues in SUMO-mediated localization of αKNL2. Furthermore, the expression of SUMO/SIM-deficient αKNL2 mutant variant in *Arabidopsis* wild type causes developmental defects, mitotic abnormalities, and impaired fertility. Nevertheless, pollen viability in the mutants remained unaffected, suggesting that meiosis is largely intact. Nevertheless, the observed fertility defects may stem from mitotic abnormalities occurring during gametophyte development or early embryogenesis. These results underscore the importance of post-translational modifications in regulating αKNL2 function in chromosome segregation and plant development. Previous studies have also shown that SUMOylation is crucial for the proper function of many centromeric proteins, and disruption of SUMOylation through mutations at SUMO attachment sites can impair these processes, causing aberrant localization. For instance, in yeast, the assembly of CENP-A/Cse4 is stimulated by C-terminal SUMO, and mutations that disrupt SUMOylation of Cse4 can lead to its mislocalization from centromeres, impacting kinetochore assembly and chromosome segregation (Ohkuni *et al*., 2018, Ohkuni *et al*., 2020). SUMOylation-deficient mutants of BubR1 show mislocalization and fail to function properly in spindle checkpoint signaling, leading to defects in chromosome segregation (Yang *et al*., 2012). In human cells, CENP-E was shown to be specifically modified by SUMO-2/3 and to have the ability to bind SUMO-2/3 polymeric chains, a function crucial for its localization to the kinetochore (Zhang *et al*., 2008).

The identification of ULP1d as a deSUMOylation enzyme for αKNL2 sheds light on the dynamic regulation of deSUMOylation in centromere function. In *ulp1d-2* mutants, enhanced SUMOylation of αKNL2-C-EYFP protein by SUMO1 and SUMO3 was observed, which resulted in severe developmental and mitotic defects. Notably, αKNL2-C-EYFP localization in *ulp1d-2* mutants shifted from the centromere to the nucleolus, highlighting the critical role of ULP1d-mediated deSUMOylation in facilitating centromere targeting of αKNL2. Interestingly, the localization of the SUMO-mutant form of αKNL2-C^Mut-SUMO^-EYFP in *ulp1d-2* mutants remained unaffected, suggesting that the deSUMOylation sites had already been altered. Similarly, deSUMOylation is mediated by a similar family of SUMO-specific proteases, including ULP in yeast and SENP in mammals, essential for centromere function. In humans, SENP6 regulates a network of proteins, including the CCAN, CENP-A loading factors Mis18BP1 and Mis18A, and DNA damage response factors, with SENP6 deficiency leading to impaired proliferation, G2/M accumulation and frequent micronuclei formation (Liebelt *et al*., 2019). In *Saccharomyces cerevisiae,* Ulp2 recruits to the kinetochore via the Ctf3^CENP-I^-Mcm16^CENP-H^-Mcm22^CENP-K^ (CMM) complex and selectively targets CCAN subunits, with mutations impairing recruitment or SUMO binding leading to increased chromosome loss and hypersensitivity to replication stress (Suhandynata *et al*., 2019).

Our study demonstrates that αKNL2 SUMOylation is essential for its association with CENH3. The interaction of αKNL2 and CENH3 was validated through BiFC and Co-IP assays; however, the yeast two-hybrid assay suggests that it may not be direct. The absence of interaction in Y2H, despite being detected in BiFC and Co-IP, indicates that αKNL2-C may require additional factors or modifications, such as SUMOylation, that are not present in yeast. Given the necessity of SUMOylation for this interaction, it is possible that this modification influences the conformation of αKNL2-C or facilitates its recruitment by other proteins. Notably, the C-terminal region of αKNL2 in Arabidopsis interacts with CENH3, as demonstrated by pull-down and BiFC assays. Consistently, structural studies in chicken show that ggKNL2, which contains a CENPC-k motif at its C-terminus, specifically recognizes CENP-A/CENH3 nucleosomes through the RG-loop in its C-terminal region (Jiang *et al*., 2023). In *Caenorhabditis elegans*, the extended N-terminal tail of CENP-A directly interacts with KNL-2, playing a crucial role in chromatin assembly and partially compensating for Scm3/HJURP function (de Groot *et al*., 2021). These findings suggest that the binding regions may vary across species, necessitating validation in each specific organism.

In plants, the SUMOylation-deficient αKNL2 mutant failed to interact with CENH3, as evidenced by both BiFC and Co-IP assays. Previous studies have shown that CENP-C interacts with CENP-A-containing nucleosomes upon CDK1-mediated phosphorylation of CENP-C in human and chicken mitotic cells (Watanabe *et al*., 2019, Ariyoshi *et al*., 2021), emphasizing the importance of PTMs in CENPA/CENH3-kinetochore interactions. Moreover, our analysis of SUMOylation-deficient αKNL2 mutants revealed a marked reduction in monomeric endogenous CENH3 and αKNL2 protein levels. In wild-type plants, CENH3 predominantly migrates as a ∼54 kDa complex, likely reflecting its incorporation into stable nucleosomal structures with associated histones. However, in the SUMOylation-deficient αKNL2 mutants, additional intermediate bands were detected, likely correspond to partially disassembled CENH3–histone subcomplexes. This suggests that impaired SUMOylation disrupts nucleosome integrity, leading to defective centromeric loading and compromised chromatin stability. These findings suggest that SUMOylation serves as a regulatory mechanism for the αKNL2-CENH3 interaction and CENH3 loading in *Arabidopsis*, which is crucial for centromere assembly and function. Our study reveals that dynamic SUMOylation of αKNL2, regulated by ULP1d, is crucial for its centromeric localization, interaction with CENH3, and role in plant development and mitosis, offering insights into centromere organization, and genome stability.

## Methods

### Plasmid construction

The complete open reading frames of SUMO1, SUMO2, SUMO3, SUMO5, and ULP1d fragments were amplified via RT-PCR using 1 µg of RNA extracted from flower buds of *A. thaliana* (Columbia-0 ecotype) wild-type, with primers listed in Supplementary Table 1. The amplified fragments were cloned into the pDONR221 backbone using the Gateway BP reaction (Invitrogen). Previously, the αKNL2, αKNL2-N, and αKNL2-C clones in the pDONR221 vector were generated as described (Lermontova et al., 2013). Fragments from the pDONR221 clones were subsequently recombined into Gateway-compatible destination vectors for various applications, such as the pGWB641 vector containing the 35S promoter was used for in vivo subcellular localization studies, while the 3’Venus-N and 3’Venus-C Gateway vectors, which include cMYC and HA tags containing the 35S promoter, were utilized for bimolecular fluorescence complementation (BiFC) and co-immunoprecipitation (Co-IP) analyses. For the negative control in co-immunoprecipitation (co-IP) experiments, an HA tag with Venus-C fragment was cloned into the 3’ Venus-C expression vector.

αKNL2-C fragments mutated simultaneously at three predicted SUMOylation sites and two predicted SUMO interaction motifs were synthesized and cloned into a Gateway-compatible pENTR-TOPO by Twist Bioscience (https://www.twistbioscience.com/). Clones with individually deleted SUMOylation sites or SUMO interaction motifs were generated from the αKNL2-C/pDONR221clone using a site-directed mutagenesis protocol (Phusion™ Site-Directed Mutagenesis Kit, Thermoscientific). Mutagenized αKNL2 fragments were recombined from pDONR221 or pENTR-TOPO clones into the Gateway-compatible pGWB641 vector via Gateway LR reaction (Invitrogen).

The C-terminal fragments of αKNL2 (pDONR221) and αKNL2^Mut-SUMO^ variant (pTwist-ENTR) were used as templates for cloning into an expression vector suitable for the SUMO *in vitro* assay. Individual fragments were amplified using specific primers listed in Supplementary Table 1. The PCR products were cloned into the BamHI site of the pET-Duet vector, which includes a His-tag at the N-terminus and a FLAG-tag at the C-terminus (Adamus *et al*., 2020), using the NEBuilder HiFi DNA Assembly Kit (New England BioLabs, USA). The cDNA from *A. thaliana* was used as a template for cloning NSE2 into the expression vector pET28c+. The PCR product was amplified using specific primers listed in Supplementary Table 1 and subsequently cloned into the BamHI and XhoI sites of the pET28c+ vector, which includes the His-tag and T7 tag at the N-terminus.

### Plant transformation and cultivation

Transient transformation of *N. benthamiana* was carried out using *Agrobacterium tumefaciens* following the protocol described by (Walter *et al*., 2004). Fluorescence signals were analyzed in the lower epidermal cell layers of tobacco leaves 48 hours post-infiltration. Each expression plasmid was introduced into *N. benthamiana* through at least three independent infiltration experiments.

For stable transformation of *A. thaliana*, plasmids were transferred into *A. tumefaciens* strain GV3101 via electroporation. Transformation of *A. thaliana* (Col-0) was performed using the floral dip method (Clough and Bent, 1998). Transgenic lines were subsequently generated, and the morphology and GFP signals resulting from plasmid-mediated gene expression were analyzed in at least three independent single-insertion lines.

Plants of *A. thaliana* and *N. benthamiana* used for localization studies, nuclei extraction, and BiFC analysis were grown under following temperature conditions. *A. thaliana* was cultivated under a 16 h light/8 h dark photoperiod with day/night temperatures of 20 °C/18 °C, while *N. benthamiana* was grown under a 12 h photoperiod at a constant temperature of 26 °C.

### Bimolecular fluorescence complementation (BiFC) assay

To visualize protein interactions in vivo, bimolecular fluorescence complementation (BiFC) was performed following the protocol of (Yadala *et al*., 2022). Leaves of 2 to 4-week-old *N. benthamiana* plants were co-infiltrated with *A. tumefaciens* strain GV3101 containing two BiFC vectors expressing the proteins of interest. HC-Pro, a RNA silencing suppressor, was co-expressed to enhance transient expression (Kasschau *et al*., 2003). Each BiFC combination was tested in at least three independent infiltration experiments using three different plants (n = 3, biological replicates).

### Yeast-two hybrid (Y2H) co-transformation assay

Yeast two-hybrid co-transformation was performed to assess the interaction between two selected proteins. The cDNAs encoding the proteins of interest were cloned into pGADT7 and pGBKT7 vectors and co-transformed into *Saccharomyces cerevisiae* Y2H Gold strain, following the Matchmaker Gold Y2H System protocol (Takara Bio, Cat #630489). Transformed yeast cells were plated on –LT (SD/-Leu/-Trp) selective media and incubated at 30 °C for 3 to 5 days. Colonies were subsequently picked, serially diluted (1/10, 1/100, 1/1000), and spotted onto –LT and –LTH (SD/-Leu/-Trp/-His) plates to evaluate interaction-dependent growth. All co-transformations and interaction assays were performed in at least two independent experiments.

### Protein extraction, immunoprecipitation (IP), and immunoblot (IB) analyses

Total protein extracts were obtained from *Arabidopsis* seedlings expressing αKNL2-C or its SUMOylation site mutants fused to EYFP using a previously described phenol extraction method (Hurkman and Tanaka, 1986). The nuclear protein extracts were isolated according to Huang et al. (2021). Protein extracts were isolated in triplicate for each genotype or treatment across 3 independent experiments. Immunoprecipitation (IP) and co-immunoprecipitation (Co-IP) were conducted according to the manufacturer’s instructions using a GFP- and HA-trap kit (Chromotek). The samples were further used for immunoblotting analyses.

For immunoblot analyses, 2X SDS sample buffer (125 mM Tris-HCl (pH 6.8), 4% SDS, 20% Glycerol, 10% β-mercaptoethanol, 0.02% Bromophenol Blue) was added to each protein sample, followed by boiling for 10 min. The protein samples were separated by SDS-PAGE using a 10% acrylamide gel and transferred to a PVDF membrane (Thermo Scientific) via electroblotting. The membranes were blocked for 1 h at room temperature in phosphate-buffered saline (PBS) containing 5% (w/v) low-fat milk powder. Membranes were then incubated overnight at 4 °C with primary antibodies diluted in 1% BSA/PBS: rabbit anti-SUMO1 (1:1000, Abcam), rabbit anti-SUMO3 (1:1000, Abcam), mouse anti-GFP (1:1000, JL8, Living Colors), mouse anti-HA (1:10,000; Proteintech), mouse anti-cMYC (1:500), mouse anti-tubulin (1:1000, Sigma, #T9026), rabbit anti-CENH3 (1:1000; Abcam, ab72001), or rabbit anti-αKNL2 (1:1000, Lifetein, rb115). Following this, the membranes were treated with secondary anti-mouse or anti-rabbit antibodies (1:5000) conjugated to IRDye 800CW and visualized using an LI-COR Odyssey scanner. The signal intensities were quantified with the Image Studio software (Version 3.1, LI-COR Biosciences).

### Protein purification

Expression vectors coding for the enzymes of the SUMO machinery (His-SUMO1 (AT4G26840), His-SUMO3 (AT5G55170), untagged SUMO-conjugating enzyme SCE1 (AT3G57870), SUMO-activating enzyme (SAE) consisting of the smaller His-tagged subunit SAE1b (AT5G50680) and the larger subunit SAE2 (AT2G21470)) were kindly provided by the laboratory of Prof. Andreas Bachmair (Tomanov *et al*., 2022).

The expression and purification of Arabidopsis SUMOylation enzymes with His-tag and αKNL2 proteins (wild-type and Mut-SUMO variants) were expressed in *Escherichia coli* BL21(DE3)RIL cells. Transformed cells were grown in LB medium at 37 °C to an optical density of 0.5 at 600 nm (OD600). Protein expression was induced with 1 mM IPTG at 37 °C for three hours. Cells were pelleted and resuspended in lysis/binding buffer (50 mM phosphate buffer, 300 mM NaCl, 10 mM imidazole, 10% glycerol, 0.5% Triton X-100, pH 8.0), followed by sonication. The lysate was cleared by centrifugation, and the supernatant was incubated with anti-His tag TALON affinity resin (Clonech, USA) for 1.5 h at 4 °C. The resin was applied to a gravity-flow column, washed with wash buffer (50 mM phosphate buffer, 300 mM NaCl, 20 mM imidazole, pH 8.0), and eluted with elution buffer containing 250 mM imidazole. Elution fractions were analyzed using SDS-PAGE and Coomassie staining. Fractions containing the target protein were pooled and concentrated using Amicon Ultra Centrifugal Filter Units (MWCO; Merck Millipore, USA). The concentrated fractions were aliquoted, and protein concentration was determined by SDS-PAGE with a BSA standard.

The NSE2 SUMO-E3 ligase was grown in LB medium at 37°C until OD600 reached 0.5. Expression was induced by 0.5mM IPTG at 30°C for 3 h. Cells were pelleted and resuspended in the same lysis/binding buffer as in the previous step with the addition of 0.5 mM TCEP. The remaining purification steps were identical to those described above. For elution, 350 mM imidazole was used, as better yields were observed under these conditions. The untagged SCE enzyme was grown in LB medium at 37°C until OD600 reached 0.5. Expression was induced by 1mM IPTG at 20°C for 1 h. After the centrifugation (4500 g, 4°C, 20 min), the pellet was resuspended in SUMO buffer (20 mM Tris–HCl, 5 mM MgCl2, 0.5 mM TCEP, pH 7.4). Lysate was sonicated, and the concentration of protein was determined using SDS-PAGE with the BSA standard.

### SUMO in vitro assay

For the SUMO in vitro assay, a similar protocol to that reported in (Tomanov *et al*., 2022) was used. The SUMOylation reaction mixture contained 2 μM SAE, 1.75 μM SCE1, 14 μM SUMO3 or SUMO1, 2 μM αKNL2 protein, 7 μM NSE2 SUMO-E3 ligase, 10× SUMO buffer, and 5 mM ATP. The reaction volume was adjusted to 20 μl with water. The mixture was incubated at 30 °C for 2 h. After incubation, 20 μl of 2x Laemmli sample buffer was added, and samples were heated at 95 °C for 5 min. Samples (20 μl) were separated on 12% SDS-PAGE gels, transferred to nitrocellulose membranes, reversibly stained with Ponceau S red, and analyzed by immunoblotting with an anti-FLAG tag HRP-conjugated antibody (Abcam A8592, 1:3000).

### Mass spectrometry (MS) analysis of in vitro SUMOylated αKNL2

For MS-based identification of SUMOylation sites, the same enzymatic system as described above for the SUMO in vitro assay was used, but scaled up to a final volume of 100 μL and SUMO proteins were substituted with cleavable H/K variants. Following the 2-hour incubation at 30 °C, the entire reaction mixture was added to 30 μL of equilibrated anti-FLAG magnetic beads (Sigma-Aldrich) and incubated at 4 °C with gentle rotation. The beads were then washed several times with buffer lacking imidazole to remove unbound proteins and contaminants. Following IP washes, bead bound protein complexes were reduced using 25mM dithiothreitol (DTT) (30 min at 56 °C) and alkylated using 100mM iodacetamide (20 min incubation at laboratory temperature in dark) with alkylation quenching done using 75mM DTT (20 min at laboratory temperature) and digested directly on beads in 50mM NaHCO3 buffer using LysC (0.2 µg, Promega) for 2h at 37 °C. Beads were removed from the initial digest and 2^nd^ digestion step took place using trypsin for 18h at 37 °C (0.5 µg; sequencing grade, Promega). Resulting peptides were extracted into LC-MS vials by 2.5% formic acid (FA) in 50% acetonitrile (ACN) and 100% ACN with addition of n-Dodecyl β-D-maltoside (DDM, final concentration 0.1%; Sigma-Aldrich) and concentrated in a SpeedVac concentrator (Thermo Fisher Scientific).

LC-MS/MS analyses of peptides solutions were done using UltiMate 3000 RSLCnano system (Thermo Fisher Scientific) connected to timsTOF Pro and or Pro 2 spectrometer (Bruker). Prior to LC separation, tryptic digests were online concentrated and desalted using a trapping column (Acclaim™ PepMap™ 100 C18, dimensions 300 μm ID, 5 mm long, 5 μm particles, Thermo Fisher Scientific). After washing the trapping column with 0.1% trifluoroacetic acid, the peptides were eluted (flow rate -150 nl/min) from the trapping column onto an analytical column (Aurora C18, 75μm ID, 250 mm long, 1.7 μm particles, heated to 50°C, PN AUR3-25075C18-CSI, Ion Opticks) by 60 min linear gradient program (3-42% of mobile phase B; mobile phase A: 0.1% FA in water; mobile phase B: 0.1% FA in 80% ACN). Equilibration of the trapping column and the analytical column was done prior to sample injection into the sample loop. The analytical column was placed inside Column Toaster heater (Bruker) and its emitter side was installed inside the CaptiveSpray ion source (Bruker) according to the manufacturer’s instructions with the column temperature set to 50 °C and spray voltage 1.4kV was used. PASEF data denoising was switched off. MS data were acquired in m/z range of 100-1700 and 1/k0 range of 0.6-1.4 V×s×cm^-2^ using DDA-PASEF method acquiring 10 PASEF scans with scheduled target intensity of 20,000 and intensity threshold of 2500. Active exclusion was set for 0.4min with precursor reconsideration for 4× more intense precursors.

Raw MS data were processed using DataAnalysis software (version 6.1) from which MS2 spectra were exported into the MGF spectra format. MS2 spectra in MGF format were database searched using Proteome Discoverer (version 1.4, Thermo Fisher Scientific) and in-house Mascot server (version 2.6.2) in the following two steps search. At first, the data were searched against modified cRAP database based on https://www.thegpm.org/crap/ using the following search parameters: precursor and fragment tolerance of 10 ppm and 0.03 Da, respectively, full tryptic/P specificity with 2 allowed missed cleavage sites, Oxidation (M), Deamidated (NQ) and Acetyl (Protein N-term) set as variable and Carbamidomethyl (C) as fixed modification. Peptide cutoff score for the cRAP database search was set to 30. Following the cRAP database search, the MS2 spectra assigned to any protein in the cRAP database with Mascot ion score 30 and above were removed and the remaining spectra were searched against specific database containing proteins of interest. Search conditions were set identically to the cRAP database search with the following modifications: additional variable modification QTGG (K), maximum number of missed cleavages 3, peptide cut off score 20. Database search results were inspected manually in Proteome Discoverer and DataAnalysis (extracted ion chromatograms) considering all acquired data inspecting and especially the following characteristics: fragments assignment quality, reported fragments delta mass, Mascot ion score and extracted ion chromatograms of the identified QTGG (K) modified peptides. Matched spectra to the SUMOylated peptides of interest were checked in results of a parallel search done against whole Arabidopsis thaliana proteome (https://ftp.uniprot.org/pub/databases/uniprot/current_release/knowledgebase/reference_proteom es/Eukaryota/UP000006548/UP000006548_3702.fasta.gz from 2025-06-19; 27,448 protein sequences) using the same conditions as the ones used against the dedicated proteins database.

### RNA isolation and RT-qPCR analysis

Total RNA was extracted from 7-day-old seedlings using TRIzol reagent and was subsequently treated with DNase to remove genomic DNA contamination. First-strand cDNA synthesis was performed using the Genaxxon Scriptase RT cDNA synthesis kit with an oligo(dT)18 primer and 2 µg of total RNA as input. Quantitative real-time PCR (RT-qPCR) was carried out on an Applied Biosystems QuantStudio 6 Flex system using Genaxxon SYBR Green Supermix. Each transcript was analyzed in triplicate across three independent biological replicates. *ACTIN* and *UBQ* were used as internal reference genes for normalization. PCR reactions (10 μL) were set up to amplify ACTIN, UBQ, αKNL2-C, CENH3. Thermal cycling conditions included an initial denaturation at 95 °C for 5 min, followed by 40 cycles of 15 s at 95 °C, 30 s at 62 °C for annealing, and 30 s at 72 °C for elongation.

### Immunostaining

Samples were prepared from *N. benthamiana* leaves infiltrated with a plasmid expressing the protein of interest tagged with EYFP. Nuclei were extracted from infiltrated leaves (Doležel *et al*., 2007). This preparation preserved EYFP fluorescence, allowing co-localization analysis via immunostaining with *N. benthamiana*-specific CENH3. For *A. thaliana*, nuclei and chromosome preparations for mitotic analysis were performed following (Lermontova *et al*., 2006). Immunostaining of nuclei and chromosomes was conducted as outlined by (Jasencakova *et al*., 2000). Primary antibodies used included rabbit anti-CENH3 (1:1000; Lifetein, rb5558), mouse anti-GFP (1:1000; Chromotek, 5F8), and mouse anti-tubulin (1:1000; Sigma, #T9026), followed by secondary antibodies: anti-rabbit rhodamine-conjugated (1:300; Jackson Immuno Research Laboratories) and anti-mouse Alexa 488 (1:300; Jackson Immuno Research Laboratories). Counterstaining was performed with 4′,6-diamidino-2-phenylindole (DAPI) (Vector Laboratories, USA).

### Microscopy

EYFP fluorescence detection was performed on transformed *N. benthamiana* leaves and *A. thaliana* seedlings using a confocal laser scanning microscope (LSM 780, Carl Zeiss GmbH). EYFP was excited with a 488-nm laser, and fluorescence was captured using a 505–550 nm band-pass filter. Spatial super-resolution structured illumination microscopy (3D-SIM) was applied using a Plan-Apochromat 63x/1.4 oil objective of an Elyra PS.1 microscope system and the software ZENblack (Carl Zeiss GmbH) (Weisshart *et al*., 2016).

### Seed set and plant fertility evaluation

For seed set analysis, siliques were fixed in ethanol:acetic acid (9:1) overnight, then dehydrated in 70% and 90% ethanol for 1 hour each. Samples were cleared overnight at 4 °C in chloral hydrate solution (chloral hydrate:water:glycerol = 8:2:1). Seeds within siliques were counted under a binocular microscope (Carl Zeiss GmbH, Germany).

To evaluate plant fertility in αKNL2 SUMOylation mutant lines, scanning electron microscopy (SEM) was used. Fresh siliques were fixed in 4% formaldehyde in 50 mM phosphate buffer (pH 7.0) for 16 h at 8 °C. Samples were washed briefly with distilled water and dehydrated in an ascending ethanol series (30%, 50%, 70%, 90%, and 100%) twice each. Critical point drying was performed using a Quorum K850 critical point dryer (Quorum Technologies Ltd.). Dried samples were mounted on carbon adhesive discs, gold-coated using an Edwards S150B sputter coater, and imaged with a Zeiss Gemini300 scanning electron microscope (Carl Zeiss Microscopy GmbH) at 5 kV acceleration voltage. Images were saved as TIFF files.

### Bioinformatic analysis

Protein interaction networks and gene ontology analyses were performed using Cytoscape v3.8.2 (https://cytoscape.org/) and STRING (https://string-db.org/). SUMOylation sites in αKNL2 were identified using the GPS-SUMO web tool (https://sumo.biocuckoo.cn/).

### Quantification and statistical analysis

Primary root lengths, silique lengths and BiFC fluorescence intensity were quantified using ImageJ software. Data are presented as mean ± SEM, based on at least three independent experiments or biological samples. For pairwise comparisons, Welch’s t-test was applied to account for unequal variances, using the T.TEST function in Excel (two-tailed, unequal variance). Comparisons involving more than two groups were analyzed by one-way ANOVA followed by Tukey’s post hoc multiple comparison test using R. Statistical significance was defined as follows: p < 0.05 (**), and p < 0.005 (***) for all tests. All statistical outputs are provided in Supplementary Data Set 1.

### Funding

This work was supported by the Deutsche Forschungsgemeinschaft DFG grants (DFG LE 2299/5-1). J.V. acknowledges support from ProteoCure COST (European Cooperation in Science and Technology) Action CA20113, which funded the research stay where J.V. acquired the methodology applied in this publication.

## Data Availability

The mass spectrometry proteomics data have been deposited to the ProteomeXchange Consortium via the PRIDE partner repository with the dataset identifier PXD051741 and 10.6019/PXD051741.

## Author Contributions

M.K. J.V. J.J.P. and I.L. conceived the study and designed experiments. M.K. J.V. V.G.J. S.D.K.Ö. performed plasmid construction, cloning, mutagenesis, localization, and protein interaction studies. M.K. and D.D. responsible for Western blot analysis. M.K. and J.R.C. performed immunostaining RT-qPCR, and mitotic analysis. J.V. B.K. D.P. and J.J.P. responsible for in vitro SUMOylation assay and MS-analysis. M.K. and V.S. performed microscopy analysis. M.K. and I.L. wrote the manuscript with the help of all co-authors. I.L. supervised and received funding for the study. All authors reviewed the manuscript.

## Acknowledgments

We thank Prof. Dr. Andreas Bachmair for providing plasmids for the SUMO *in vitro* assay. Dr. Eva Dvořák Tomaštíková for providing anti-SUMO1 and anti-SUMO3 antibodies. Dr. Twan Rutten for SEM microscopy analysis. Heike Kuhlmann and Annette Heber for technical assistance.

## Supplementary materials

**Supplementary Figure 1.** The post-translational modification pathway analysis of αKNL2 interactors based on Y2H screening

**Supplementary Figure 2.** The interaction analysis of SUMO and ULP1d proteins with αKNL2 by BiFC

**Supplementary Figure 3.** The conservation analysis of SUMOylation and SIM sites in αKNL2-C

**Supplementary Figure 4.** Immunoblot detection of αKNL2-C-EYFP and SUMOylation-deficient αKNL2-C^Mut-SUMO^-EYFP in Arabidopsis transgenic lines.

**Supplementary Figure 5.** The interaction of SUMO3 and ULP1d with SUMO mutant variants of αKNL2-C by BiFC

**Supplementary Figure 6.** Analysis of chromosome segregation defects, pollen viability, and seed set in the SUMOylation-deficient αKNL2 mutant

**Supplementary Figure 7.** The in vivo and in vitro SUMOylation analysis of αKNL2-C

**Supplementary Figure 8.** The phenotype characteristics and localization of αKNL2-C and αKNL2-C^Mut-SUMO^ in *ulp1d-2*

**Supplementary Figure 9.** Transcript levels of *αKNL2* and SUMO3 western blot in αKNL2-C and αKNL2-C^Mut-SUMO^ lines in wild-type and *ulp1d-2* mutant plants

**Supplemental Figure 10.** BiFC quantification and yeast two-hybrid assay for interactions between αKNL2 and CENH3

**Supplementary Figure 11.** αKNL2 protein levels are reduced in the SUMOylation-deficient αKNL2-C mutant

**Supplementary Table 1.** Primers used in this study

**Supplementary Data Set 1**. Summary of statistical analysis.

## References

Adamus, M., Lelkes, E., Potesil, D., Ganji, S.R., Kolesár, P., Zábrady, K., Zdráhal, Z. and Palecek, J. (2020) Molecular insights into the architecture of the human SMC5/6 complex. Journal of molecular biology, 432, 3820–3837.

Andrews, E.A., Palecek, J., Sergeant, J., Taylor, E., Lehmann, A.R. and Watts, F.Z. (2005) Nse2, a component of the Smc5-6 complex, is a SUMO ligase required for the response to DNA damage. Molecular and cellular biology.

Aragón, L. (2018) The Smc5/6 complex: new and old functions of the enigmatic long-distance relative. Annual review of genetics, 52, 89–107.

Ariyoshi, M., Makino, F., Watanabe, R., Nakagawa, R., Kato, T., Namba, K., Arimura, Y., Fujita, R., Kurumizaka, H., Okumura, E.i., Hara, M. and Fukagawa, T. (2021) Cryo-EM structure of the CENP-A nucleosome in complex with phosphorylated CENP-C. The EMBO Journal, 40, e105671.

Azuma, Y., Arnaoutov, A. and Dasso, M. (2003) SUMO-2/3 regulates topoisomerase II in mitosis. The Journal of cell biology, 163, 477–487.

Bailey, M., Srivastava, A., Conti, L., Nelis, S., Zhang, C., Florance, H., Love, A., Milner, J., Napier, R. and Grant, M. (2016) Stability of small ubiquitin-like modifier (SUMO) proteases OVERLY TOLERANT TO SALT1 and-2 modulates salicylic acid signalling and SUMO1/2 conjugation in Arabidopsis thaliana. Journal of Experimental Botany, 67, 353–363.

Ban, R., Nishida, T. and Urano, T. (2011) Mitotic kinase Aurora-B is regulated by SUMO-2/3 conjugation/deconjugation during mitosis. Genes to Cells, 16, 652–669.

Castaño-Miquel, L., Seguí, J. and Lois, L.M. (2011) Distinctive properties of Arabidopsis SUMO paralogues support the in vivo predominant role of AtSUMO1/2 isoforms. Biochemical Journal, 436, 581–590.

Castro, P.H., Couto, D., Freitas, S., Verde, N., Macho, A.P., Huguet, S., Botella, M.A., Ruiz-Albert, J., Tavares, R.M. and Bejarano, E.R. (2016) SUMO proteases ULP1c and ULP1d are required for development and osmotic stress responses in Arabidopsis thaliana. Plant Molecular Biology, 92, 143–159.

Chosed, R., Mukherjee, S., Lois, L.M. and Orth, K. (2006) Evolution of a signalling system that incorporates both redundancy and diversity: Arabidopsis SUMOylation. Biochemical Journal, 398, 521–529.

Chupreta, S., Holmstrom, S., Subramanian, L. and Iñiguez-Lluhí, J.A. (2005) A small conserved surface in SUMO is the critical structural determinant of its transcriptional inhibitory properties. Molecular and cellular biology, 25, 4272–4282.

Clough, S.J. and Bent, A.F. (1998) Floral dip: a simplified method for Agrobacterium-mediated transformation of Arabidopsis thaliana. The plant journal, 16, 735–743.

Colby, T., Matthäi, A., Boeckelmann, A. and Stuible, H.-P. (2006) SUMO-conjugating and SUMO-deconjugating enzymes from Arabidopsis. Plant Physiology, 142, 318–332.

Conti, L., Nelis, S., Zhang, C., Woodcock, A., Swarup, R., Galbiati, M., Tonelli, C., Napier, R., Hedden, P. and Bennett, M. (2014) Small ubiquitin-like modifier protein SUMO enables plants to control growth independently of the phytohormone gibberellin. Developmental cell, 28, 102–110.

Conti, L., Price, G., O’Donnell, E., Schwessinger, B., Dominy, P. and Sadanandom, A. (2008) Small ubiquitin-like modifier proteases OVERLY TOLERANT TO SALT1 and-2 regulate salt stress responses in Arabidopsis. The Plant Cell, 20, 2894–2908.

Cubeñas-Potts, C., Goeres, J.D. and Matunis, M.J. (2013) SENP1 and SENP2 affect spatial and temporal control of sumoylation in mitosis. Molecular biology of the cell, 24, 3483–3495.

Cuijpers, S.A., Willemstein, E. and Vertegaal, A.C. (2017) Converging small ubiquitin-like modifier (SUMO) and ubiquitin signaling: improved methodology identifies co-modified target proteins. Molecular & Cellular Proteomics, 16, 2281–2295.

de Groot, C., Houston, J., Davis, B., Gerson-Gurwitz, A., Monen, J., Lara-Gonzalez, P., Oegema, K., Shiau, A.K. and Desai, A. (2021) The N-terminal tail of C. elegans CENP-A interacts with KNL-2 and is essential for centromeric chromatin assembly. Molecular biology of the cell, 32, 1193–1201.

Doležel, J., Greilhuber, J. and Suda, J. (2007) Estimation of nuclear DNA content in plants using flow cytometry. Nature protocols, 2, 2233–2244.

Fernández-Miranda, G., de Castro, I.P., Carmena, M., Aguirre-Portolés, C., Ruchaud, S., Fant, X., Montoya, G., Earnshaw, W.C. and Malumbres, M. (2010) SUMOylation modulates the function of Aurora-B kinase. Journal of cell science, 123, 2823–2833.

French, B.T., Westhorpe, F.G., Limouse, C. and Straight, A.F. (2017) *Xenopus laevis* M18BP1 directly binds existing CENP-A nucleosomes to promote centromeric chromatin assembly. Dev Cell., 42, 190–199 e110.

Fu, H., Liu, N., Dong, Q., Ma, C., Yang, J., Xiong, J., Zhang, Z., Qi, X., Huang, C. and Zhu, B. (2019) SENP6-mediated M18BP1 deSUMOylation regulates CENP-A centromeric localization. Cell Research, 29, 254–257.

Gong, L. and Yeh, E.T. (2006) Characterization of a family of nucleolar SUMO-specific proteases with preference for SUMO-2 or SUMO-3. Journal of Biological Chemistry, 281, 15869–15877.

Hay, R.T. (2005) SUMO: a history of modification. Molecular cell, 18, 1–12.

Hermkes, R., Fu, Y.-F., Nürrenberg, K., Budhiraja, R., Schmelzer, E., Elrouby, N., Dohmen, R.J., Bachmair, A. and Coupland, G. (2011) Distinct roles for Arabidopsis SUMO protease ESD4 and its closest homolog ELS1. Planta, 233, 63–73.

Hori, T., Shang, W.H., Hara, M., Ariyoshi, M., Arimura, Y., Fujita, R., Kurumizaka, H. and Fukagawa, T. (2017) Association of M18BP1/KNL2 with CENP-A Nucleosome Is Essential for Centromere Formation in Non-mammalian Vertebrates. Dev Cell, 42, 181–189 e183.

Hurkman, W.J. and Tanaka, C.K. (1986) Solubilization of plant membrane proteins for analysis by two-dimensional gel electrophoresis. Plant physiology, 81, 802–806.

Jasencakova, Z., Meister, A., Walter, J., Turner, B.M. and Schubert, I. (2000) Histone H4 acetylation of euchromatin and heterochromatin is cell cycle dependent and correlated with replication rather than with transcription. The Plant Cell, 12, 2087–2100.

Jiang, H., Ariyoshi, M., Hori, T., Watanabe, R., Makino, F., Namba, K. and Fukagawa, T. (2023) The cryo-EM structure of the CENP-A nucleosome in complex with ggKNL2. The EMBO Journal, 42, e111965.

Kalidass, M., Jarubula, V.G., Ratnikava, M., Chandra, J.R., Le Goff, S., Probst, A.V., Esposito, S., Grasser, K.D., Bruckmann, A. and Gagneux, J.F. (2025) Ubiquitin-dependent proteolysis of KNL2 driven by APC/CCDC20 is critical for centromere integrity and mitotic fidelity. The Plant Cell, 37, koaf164.

Kasschau, K.D., Xie, Z., Allen, E., Llave, C., Chapman, E.J., Krizan, K.A. and Carrington, J.C. (2003) P1/HC-Pro, a Viral Suppressor of RNA Silencing, Interferes with <EM>Arabidopsis</EM> Development and miRNA Function. Developmental Cell, 4, 205–217.

Lermontova, I., Kuhlmann, M., Friedel, S., Rutten, T., Heckmann, S., Sandmann, M., Demidov, D., Schubert, V. and Schubert, I. (2013) Arabidopsis kinetochore null2 is an upstream component for centromeric histone H3 variant cenH3 deposition at centromeres. The Plant Cell, 25, 3389–3404.

Lermontova, I., Schubert, V., Fuchs, J., Klatte, S., Macas, J. and Schubert, I. (2006) Loading of Arabidopsis centromeric histone CENH3 occurs mainly during G2 and requires the presence of the histone fold domain. The Plant Cell, 18, 2443–2451.

Li, T., Chen, L., Cheng, J., Dai, J., Huang, Y., Zhang, J., Liu, Z., Li, A., Li, N. and Wang, H. (2016) SUMOylated NKAP is essential for chromosome alignment by anchoring CENP-E to kinetochores. Nature communications, 7, 12969.

Liebelt, F., Jansen, N.S., Kumar, S., Gracheva, E., Claessens, L.A., Verlaan-de Vries, M., Willemstein, E. and Vertegaal, A.C. (2019) The poly-SUMO2/3 protease SENP6 enables assembly of the constitutive centromere-associated network by group deSUMOylation. Nature Communications, 10, 3987.

Mahajan, R., Gerace, L. and Melchior, F. (1998) Molecular characterization of the SUMO-1 modification of RanGAP1 and its role in nuclear envelope association. The Journal of cell biology, 140, 259–270.

Matunis, M.J., Wu, J. and Blobel, G. (1998) SUMO-1 modification and its role in targeting the Ran GTPase-activating protein, RanGAP1, to the nuclear pore complex. The Journal of cell biology, 140, 499–509.

Mérai, Z., Chumak, N., García-Aguilar, M., Hsieh, T.-F., Nishimura, T., Schoft, V.K., Bindics, J., Ślusarz, L., Arnoux, S. and Opravil, S. (2014) The AAA-ATPase molecular chaperone Cdc48/p97 disassembles sumoylated centromeres, decondenses heterochromatin, and activates ribosomal RNA genes. Proceedings of the National Academy of Sciences, 111, 16166–16171.

Miller, K.E., Kim, Y., Huh, W.-K. and Park, H.-O. (2015) Bimolecular fluorescence complementation (BiFC) analysis: advances and recent applications for genome-wide interaction studies. Journal of molecular biology, 427, 2039–2055.

Montpetit, B., Hazbun, T.R., Fields, S. and Hieter, P. (2006) Sumoylation of the budding yeast kinetochore protein Ndc10 is required for Ndc10 spindle localization and regulation of anaphase spindle elongation. The Journal of cell biology, 174, 653–663.

Mukhopadhyay, D., Arnaoutov, A. and Dasso, M. (2010) The SUMO protease SENP6 is essential for inner kinetochore assembly. Journal of Cell Biology, 188, 681–692.

Müller, S., Hoege, C., Pyrowolakis, G. and Jentsch, S. (2001) SUMO, ubiquitin’s mysterious cousin. Nature reviews Molecular cell biology, 2, 202–210.

Murtas, G., Reeves, P.H., Fu, Y.-F., Bancroft, I., Dean, C. and Coupland, G. (2003) A nuclear protease required for flowering-time regulation in Arabidopsis reduces the abundance of SMALL UBIQUITIN-RELATED MODIFIER conjugates. The Plant Cell, 15, 2308–2319.

Naish, M. and Henderson, I.R. (2024) The structure, function, and evolution of plant centromeres. Genome Research, 34, 161–178.

Ohkuni, K., Levy-Myers, R., Warren, J., Au, W.-C., Takahashi, Y., Baker, R.E. and Basrai, M.A. (2018) N-terminal sumoylation of centromeric histone H3 variant Cse4 regulates its proteolysis to prevent mislocalization to non-centromeric chromatin. G3: Genes, Genomes, Genetics, 8, 1215–1223.

Ohkuni, K., Suva, E., Au, W.-C., Walker, R.L., Levy-Myers, R., Meltzer, P.S., Baker, R.E. and Basrai, M.A. (2020) Deposition of centromeric histone H3 variant CENP-A/Cse4 into chromatin is facilitated by its C-terminal sumoylation. Genetics, 214, 839–854.

Palecek, J.J. (2018) SMC5/6: multifunctional player in replication. Genes, 10, 7.

Park, H.J., Kim, W.-Y., Park, H.C., Lee, S.Y., Bohnert, H.J. and Yun, D.-J. (2011) SUMO and SUMOylation in plants. Molecules and cells, 32, 305–316.

Pichler, A., Knipscheer, P., Oberhofer, E., Van Dijk, W.J., Körner, R., Olsen, J.V., Jentsch, S., Melchior, F. and Sixma, T.K. (2005) SUMO modification of the ubiquitin-conjugating enzyme E2-25K. Nature structural & molecular biology, 12, 264–269.

Roy, D. and Sadanandom, A. (2021) SUMO mediated regulation of transcription factors as a mechanism for transducing environmental cues into cellular signaling in plants. Cellular and Molecular Life Sciences, 78, 2641–2664.

Sadanandom, A., Ádám, É., Orosa, B., Viczián, A., Klose, C., Zhang, C., Josse, E.-M., Kozma-Bognár, L. and Nagy, F. (2015) SUMOylation of phytochrome-B negatively regulates light-induced signaling in Arabidopsis thaliana. Proceedings of the National Academy of Sciences, 112, 11108–11113.

Sandmann, M., Talbert, P., Demidov, D., Kuhlmann, M., Rutten, T., Conrad, U. and Lermontova, I. (2017) Targeting of Arabidopsis KNL2 to Centromeres Depends on the Conserved CENPC-k Motif in Its C Terminus. Plant Cell, 29, 144–155.

Subramonian, D., Chen, T.-A., Paolini, N. and Zhang, X.-D.D. (2021) Poly-SUMO-2/3 chain modification of Nuf2 facilitates CENP-E kinetochore localization and chromosome congression during mitosis. Cell Cycle, 20, 855–873.

Suhandynata, R.T., Quan, Y., Yang, Y., Yuan, W.-T., Albuquerque, C.P. and Zhou, H. (2019) Recruitment of the Ulp2 protease to the inner kinetochore prevents its hyper-sumoylation to ensure accurate chromosome segregation. PLoS genetics, 15, e1008477.

Talbert, P.B., Masuelli, R., Tyagi, A.P., Comai, L. and Henikoff, S. (2002) Centromeric localization and adaptive evolution of an Arabidopsis histone H3 variant. The Plant Cell, 14, 1053–1066.

Tomanov, K., Julian, J., Ziba, I. and Bachmair, A. (2022) SUMO Conjugation and SUMO Chain Formation by Plant Enzymes. In Plant Proteostasis: Methods and Protocols: Springer, pp. 83–92.

van den Berg, S.J., East, S., Mitra, S. and Jansen, L.E. (2023) p97/VCP drives turnover of SUMOylated centromeric CCAN proteins and CENP-A. Molecular Biology of the Cell, 34, br6.

van den Berg, S.J. and Jansen, L.E. (2023) SUMO control of centromere homeostasis. Frontiers in Cell and Developmental Biology, 11, 1193192.

Walter, M., Chaban, C., Schütze, K., Batistic, O., Weckermann, K., Näke, C., Blazevic, D., Grefen, C., Schumacher, K. and Oecking, C. (2004) Visualization of protein interactions in living plant cells using bimolecular fluorescence complementation. The Plant Journal, 40, 428–438.

Watanabe, R., Hara, M., Okumura, E.-i., Hervé, S., Fachinetti, D., Ariyoshi, M. and Fukagawa, T. (2019) CDK1-mediated CENP-C phosphorylation modulates CENP-A binding and mitotic kinetochore localization. Journal of Cell Biology, 218, 4042–4062.

Weisshart, K., Fuchs, J. and Schubert, V. (2016) Structured illumination microscopy (SIM) and photoactivated localization microscopy (PALM) to analyze the abundance and distribution of RNA polymerase II molecules on flow-sorted Arabidopsis nuclei. Bio-protocol, 6, e1725–e1725.

Wilson, V.G. and Rangasamy, D. (2001) Intracellular targeting of proteins by sumoylation. Experimental cell research, 271, 57–65.

Yadala, R., Ratnikava, M. and Lermontova, I. (2022) Bimolecular Fluorescence Complementation to Test for Protein–Protein Interactions and to Uncover Regulatory Mechanisms During Gametogenesis. In Plant Gametogenesis: Methods and Protocols: Springer, pp. 107–120.

Yalagapati, S.P., Ahmadli, U., Sinha, A., Kalidass, M., Dabravolski, S., Zuo, S., Yadala, R., Rutten, T., Talbert, P. and Berr, A. (2024) Centromeric localization of αKNL2 and CENP-C proteins in plants depends on their centromere-targeting domain and DNA-binding regions. *Nucleic Acids Research*, gkae1242.

Yang, F., Hu, L., Chen, C., Yu, J., O’Connell, C.B., Khodjakov, A., Pagano, M. and Dai, W. (2012) BubR1 is modified by sumoylation during mitotic progression. Journal of Biological Chemistry, 287, 4875–4882.

Zhang, D., Martyniuk, C.J. and Trudeau, V.L. (2006) SANTA domain: a novel conserved protein module in Eukaryota with potential involvement in chromatin regulation. Bioinformatics, 22, 2459–2462.

Zhang, X.-D., Goeres, J., Zhang, H., Yen, T.J., Porter, A.C. and Matunis, M.J. (2008) SUMO-2/3 modification and binding regulate the association of CENP-E with kinetochores and progression through mitosis. Molecular cell, 29, 729–741.

