## Supplementary Figures for "The C-terminal SUMOylation-dependent regulation of αKNL2 governs its centromere targeting and interaction with CENH3"

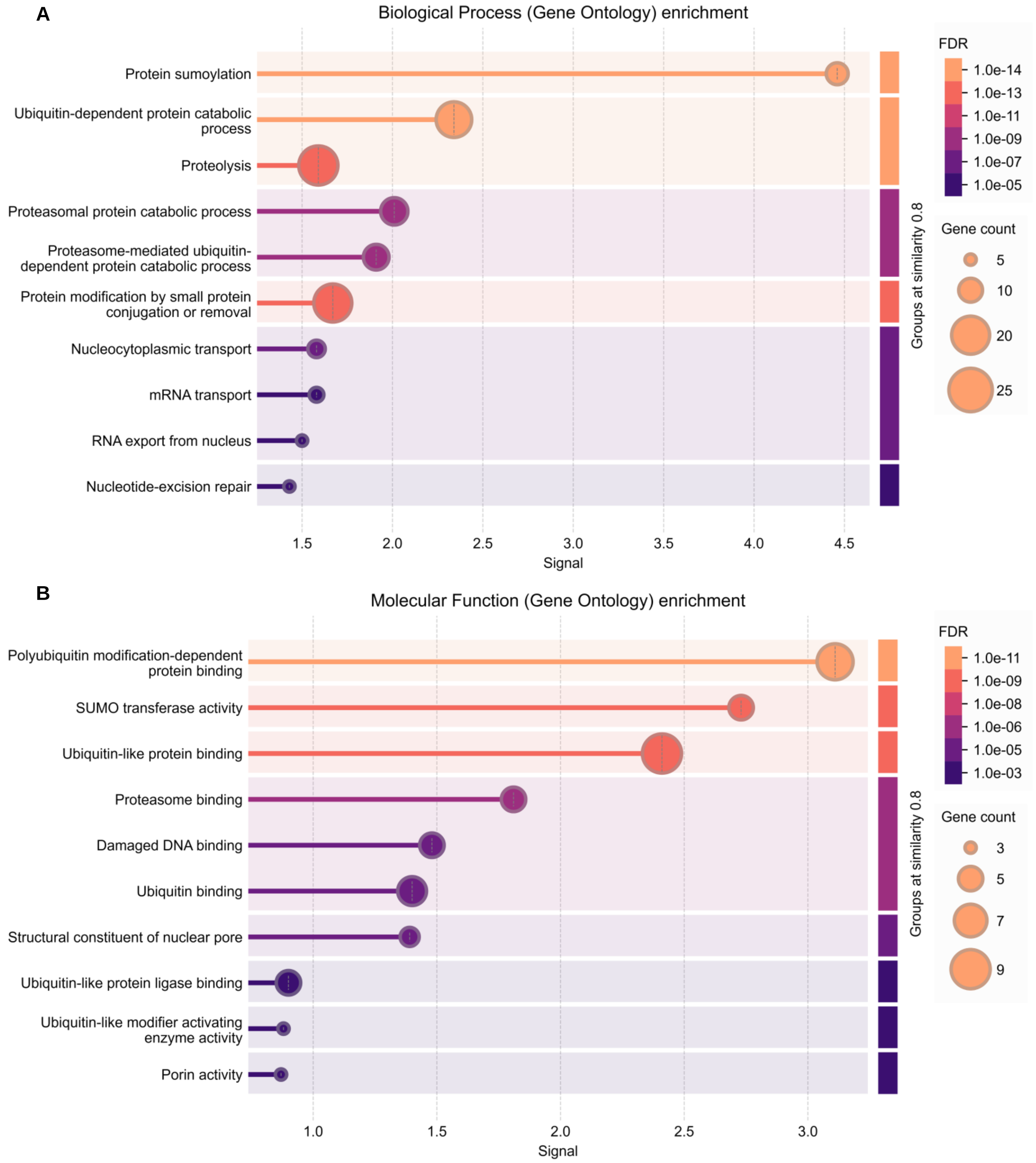


**Supplementary Figure 1. The post-translational modification pathway analysis of αKNL2 interactors based on Y2H screening**

**(A, B)** The gene ontology analysis such as biological process **(A)** and molecular function **(B)** of post-translational modification of αKNL2 interactors identified proteolysis, SUMOylation and transport pathways.

**A**

| | | C-YFP | | | | | | | |
|---|---|---|---|---|---|---|---|---|---|
| | | SUMO1 | SUMO2 | SUMO3 | SUMO5 | ULP1d | ULP1d-C | ULP1d-N | Empty |
| N-YFP | αKNL2 | | | | | | | | |
| | αKNL2-C | | | Nucleolus | | Nucleolus | Nucleolus | | |
| | αKNL2-N | | | | | | | | |
| | Empty | | | | | | | | |

**B**

| | | C-YFP | | | |
|---|---|---|---|---|---|
| | | αKNL2 | αKNL2-C | αKNL2-N | Empty |
| N-YFP | SUMO1 | | | | |
| | SUMO2 | | | | |
| | SUMO3 | | Nucleolus | | |
| | SUMO5 | | | | |
| | ULP1d | | Nucleolus | | |
| | ULP1d-C | | Nucleolus | | |
| | ULP1d-N | | | | |
| | Empty | | | | |

**Supplementary Figure 2. The interaction analysis of SUMO and ULP1d proteins with αKNL2 by BiFC**

The BiFC interactions shown for the combinations such as αKNL2, αKNL2-N, αKNL2-C fused to VENn and SUMO1, SUMO2, SUMO3, SUMO5, ULP1d, ULP1d-N or ULP1d-C fused to VENc **(A)** and vice versa **(B)**. SUMO3 and ULP1d showed interaction only with αKNL2-C, while other SUMO proteins did not interact with any αKNL2 variants. The empty BiFC negative controls were used to validate the positive interactions.

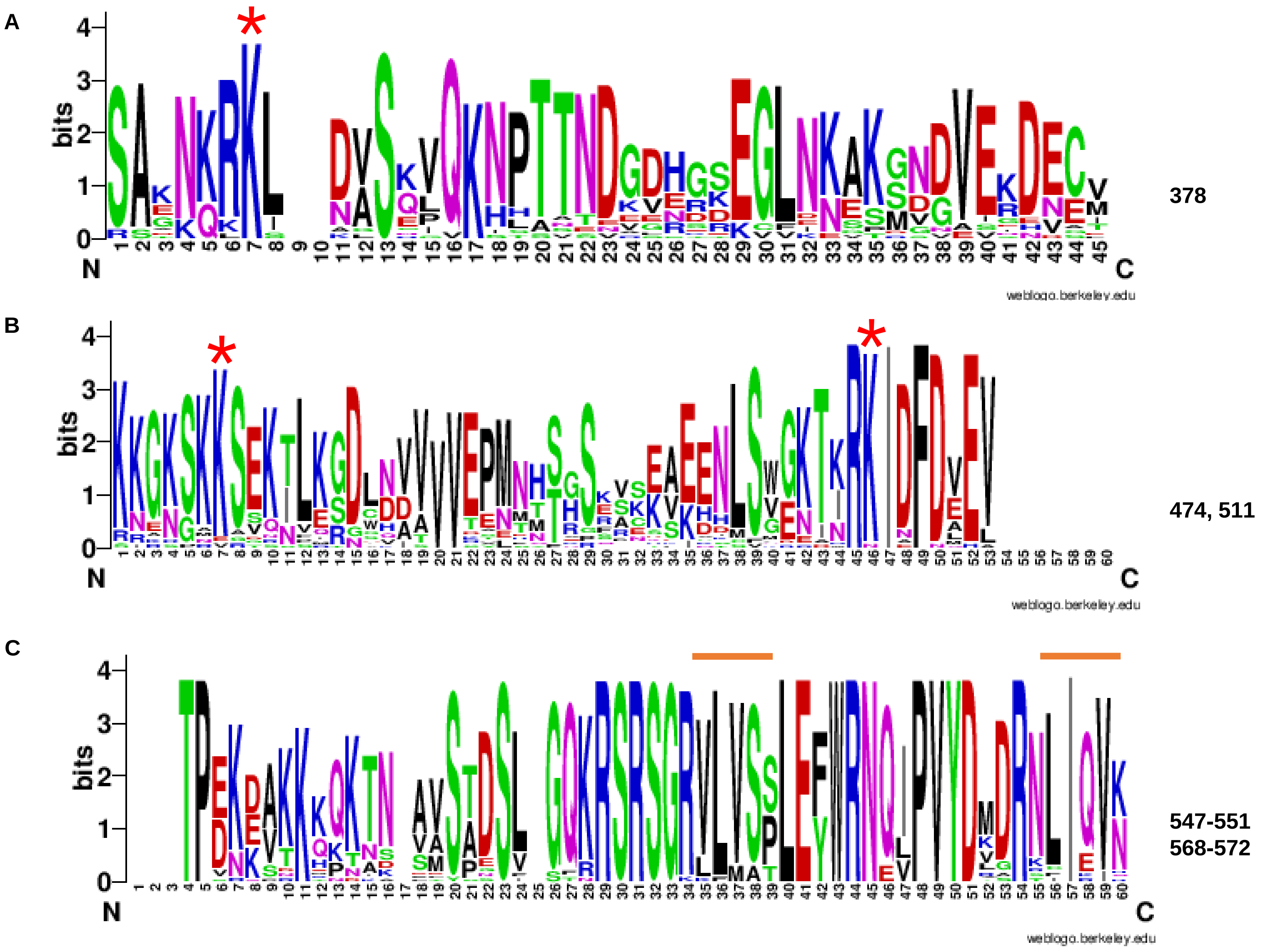


**Supplementary Figure 3. The conservation analysis of SUMOylation and SIM sites in αKNL2-C**

**(A–C)** The conservation of SUMOylation and SIM sites predicted by GPS-SUMO in αKNL2-C across Brassicales species, as illustrated using WebLogo ([https://weblogo.berkeley.edu/logo.cgi](https://weblogo.berkeley.edu/logo.cgi)). Conserved lysine residues and SUMO interaction sites are indicated by red asterisks and orange lines, respectively.

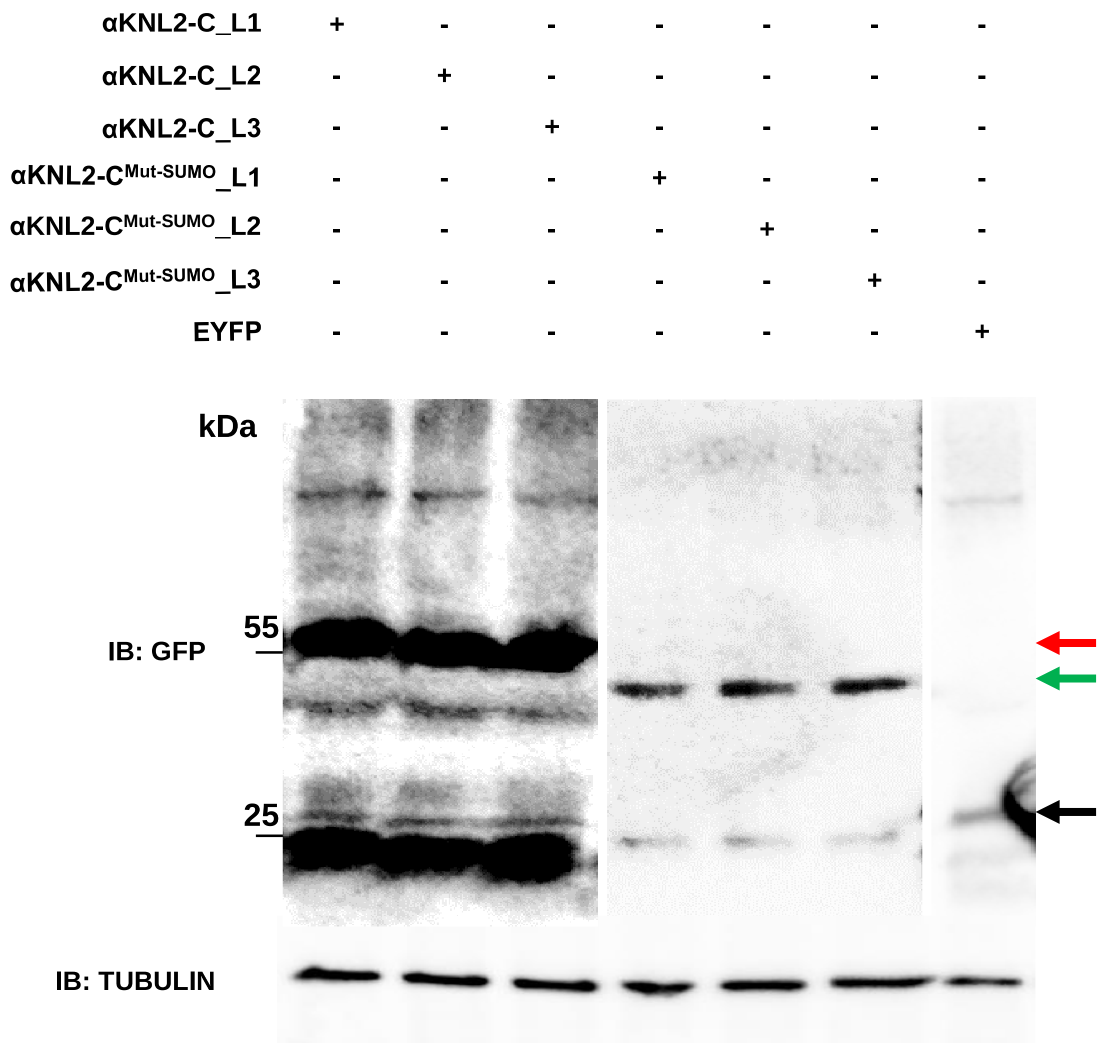


**Supplementary Figure 4. Immunoblot detection of αKNL2-C-EYFP and SUMOylation-deficient αKNL2-C$^{\text{Mut-SUMO}}$-EYFP in Arabidopsis transgenic lines.**

Total protein extracts from three independent transgenic lines expressing αKNL2-C-EYFP, SUMOylation-deficient mutant αKNL2-C$^{\text{Mut-SUMO}}$-EYFP or EYFP alone were subjected to GFP immunoblotting. The red arrow, green and black indicates the expected size of the αKNL2-C-EYFP, αKNL2-C$^{\text{Mut-SUMO}}$-EYFP and EYFP fusion protein, respectively. The tubulin was used as a loading control to confirm equal protein loading.

**A** SUMO3-VENn + αKNL2-C$^{Mut\text{-}SUMO}$-VENc

**B** ULP1d-VENn + αKNL2-C$^{Mut\text{-}SUMO}$-VENc

**C** ULP1d-C-VENn + αKNL2-C$^{Mut\text{-}SUMO}$-VENc

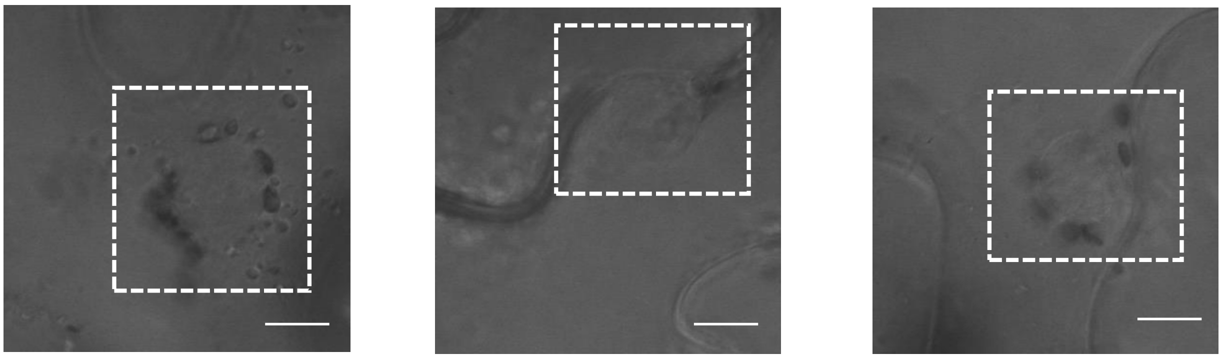

**D**

| | | VENUS-C | | |
|---|---|---|---|---|
| | Proteins | SUMO3 | ULP1d | ULP1d-C |
| VENUS-N | αKNL2 | | | |
| | αKNL2-C | Nucleolus | Nucleolus | Nucleolus |
| | αKNL2-N | | | |
| | αKNL2-C$^{Mut\text{-}SUMO}$ | | | |

Positive interaction    No interaction    No interaction after mutagenesis

**E**

| | | VENUS-C | | |
|---|---|---|---|---|
| | Proteins | SUMO3 | ULP1d | ULP1d-C |
| VENUS-N | K378R--αKNL2-C | Nucleolus | Nucleolus | Nucleolus |
| | K474R--αKNL2-C | Nucleolus | Nucleolus | Nucleolus |
| | K511R--αKNL2-C | Nucleolus | Nucleolus | Nucleolus |
| | Δ547-551--αKNL2-C | Nucleolus | Nucleolus | Nucleolus |
| | Δ568-572--αKNL2-C | Nucleolus | Nucleolus | Nucleolus |
| | K378R-K474R--αKNL2-C | Nucleolus | Nucleolus | Nucleolus |
| | K474R-K511R--αKNL2-C | | | |

Positive interaction    No interaction    No interaction after mutagenesis

**Supplementary Figure 5. The interaction of SUMO3 and ULP1d with SUMO mutant variants of αKNL2-C by BiFC**

**(A-C)** No interaction was found between SUMO3, ULP1d, ULP1d-C fused to VENn with αKNL2-C$^{Mut\text{-}SUMO}$ fused to VENc. The nucleus lacking Venus fluorescence is indicated by white dotted circles. Scale bars represents 5 µm. **(D)** The similar interaction results was found when the orientation of the Venus fusion was reversed. The interaction was indicated in a colour code below the table. **(E)** Analysis of the interactions between individual SUMOylation and SIM sites of αKNL2-C with SUMO3, ULP1d, or ULP1d-C. Double lysine mutations (K474R and K511R) abolished interactions with SUMO3 and ULP1d. Interaction outcomes are indicated by a color code below the table.

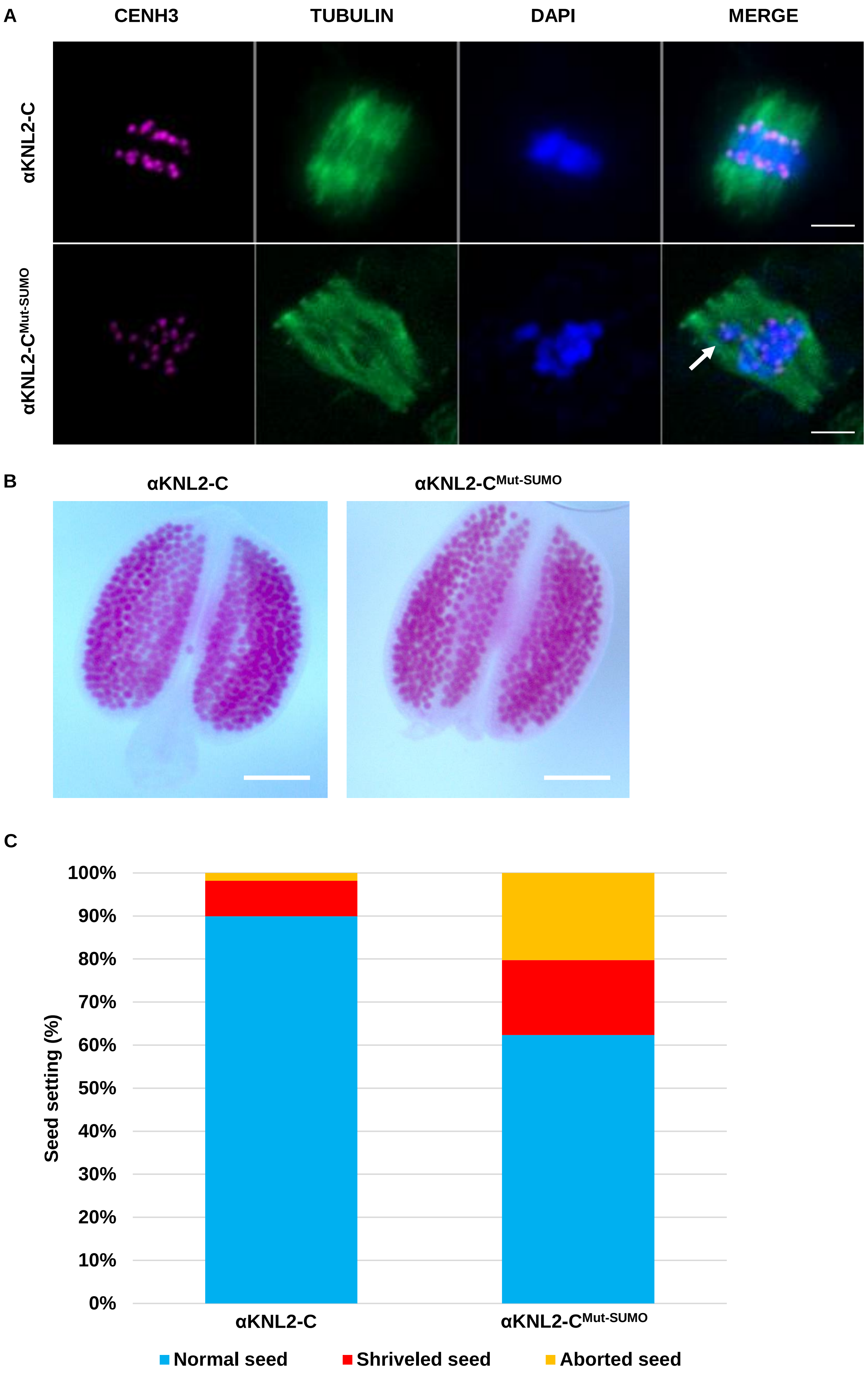


**Supplementary Figure 6. Analysis of chromosome segregation defects, pollen viability, and seed set in the SUMOylation-deficient αKNL2 mutant**

**(A)** Representative images of mitotic chromosome segregation in Arabidopsis αKNL2 SUMO mutant plants. Microtubules (green) are stained with anti-tubulin, centromeres (magenta) with anti-CENH3, and DNA (blue) with DAPI. The upper row shows normal metaphase alignment, while the lower panel depicts a misaligned chromosome (arrow) during metaphase. Scale bars 5 µm. **(B)** Alexander staining of pollen grains from αKNL2-C (left) and αKNL2$^{Mut-SUMO}$ mutant (right) plants. The mutant anthers did not show any difference compared to αKNL2-C control plants, suggesting no defects in pollen viability. Scale bars 10 µm. **(C)** Analysis of seed setting in *Arabidopsis* αKNL2-C or αKNL2-C$^{Mut-SUMO}$ mutant variant. Bar graph showing the number of normal, shriveled and aborted seeds per silique for 10 plants per construct, with 10 siliques analyzed per plant.

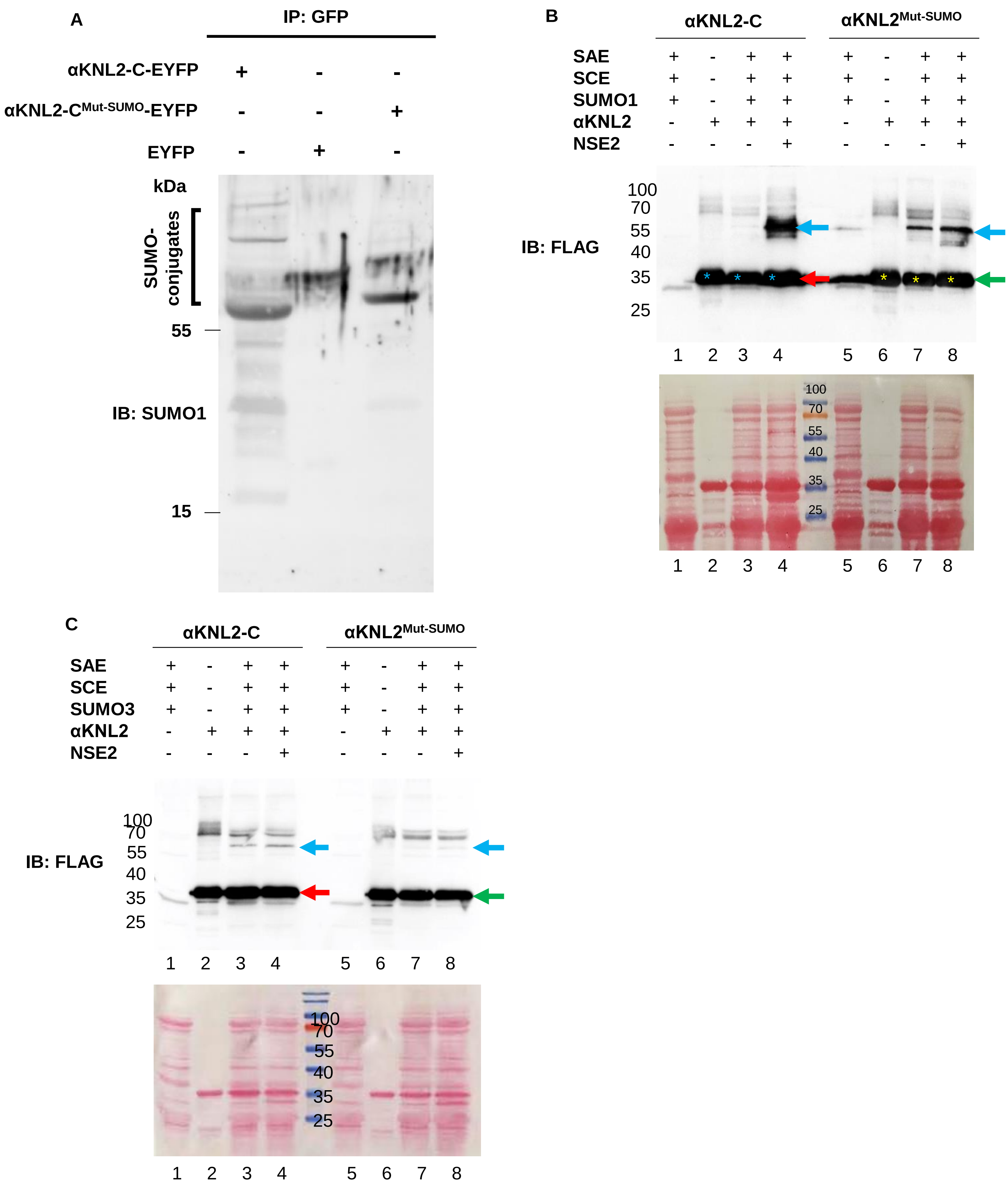


**Supplementary Figure 7. The in vivo and in vitro SUMOylation analysis of αKNL2-C**

**(A)** In vivo SUMOylation assay of αKNL2 was performed in *N. benthamiana* leaves expressing αKNL2-C-EYFP, αKNL2-C$^{Mut\text{-}SUMO}$-EYFP or EYFP alone. Total protein extracts were subjected to immunoprecipitation using GFP beads, followed by immunoblotting with anti-SUMO1 antibodies. Input controls and tubulin loading controls are presented in Fig. 5. SUMO conjugates are indicated by black brackets. Abbreviations: IB, Immunoblot; IP, Immunoprecipitation. **(B, C)** The in vitro SUMOylation assay was performed to assess the SUMOylation efficiency of αKNL2-C and its SUMO mutant variant using the SUMO1 (**B**) or SUMO3 (**C**) isoforms. The reactions included enzymes only (lanes 1 and 5), substrate only (lanes 2 and 6), a mixture of enzymes and substrate (lanes 3 and 7), and a complete reaction with the addition of NSE2 SUMO-E3 ligase (lanes 4 and 8). Following incubation, samples were analyzed using SDS-PAGE and immunoblotting. The membrane was reversibly stained with Ponceau S red (lower panels) as a loading control to verify equal protein loading across all reactions. Both αKNL2 variants and their SUMOylated forms were detected using an anti-FLAG antibody (upper panels). The red arrows represents the unmodified αKNL2-C, the green arrows marks the unmodified αKNL2-C$^{Mut\text{-}SUMO}$ mutant, and the blue arrows indicate SUMOylated forms. The SUMO mutant shows a significantly reduced SUMOylation efficiency compared to the wild-type variant. Notably, the addition of NSE2 did not enhance SUMO3 efficiency for either αKNL2 variant, while it enhanced SUMO1 efficiency for both variants.

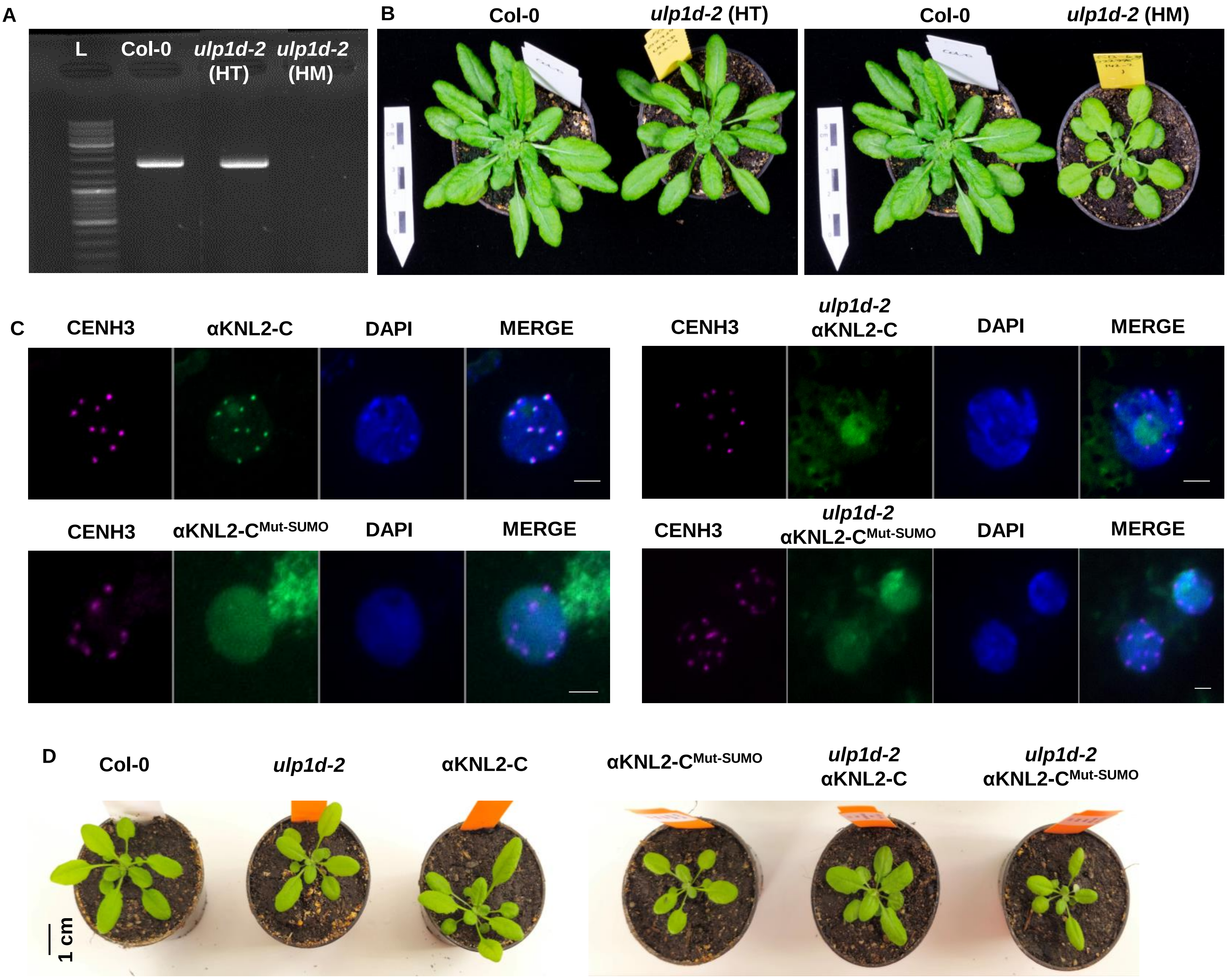


**Supplementary Figure 8. The phenotype characteristics and localization of αKNL2-C and αKNL2-C$^{Mut\text{-}SUMO}$ in *ulp1d-2***

**(A-B)** RT-PCR analysis of ULP1d expression (A) and the phenotype of 5 weeks grown plants (B) of wild-type (Col-0), *ulp1d-2* heterozygous and homozygous backgrounds. **(C)** Immunostaining experiments showing the co-localization of αKNL2-C and KNL2-C$^{Mut\text{-}SUMO}$ (green) in meristematic nuclei of wild-type and *ulp1d-2* mutants. The nuclei were stained with anti-CENH3 (red) and DAPI was used as a counterstain. Scale bars represents 5 µm. **(D)** The phenotype of the αKNL2-C and KNL2-C$^{Mut\text{-}SUMO}$ in wild-type and *ulp1d-2* mutants grown for 4 weeks in soil. Scale bar represents 1 cm.

A

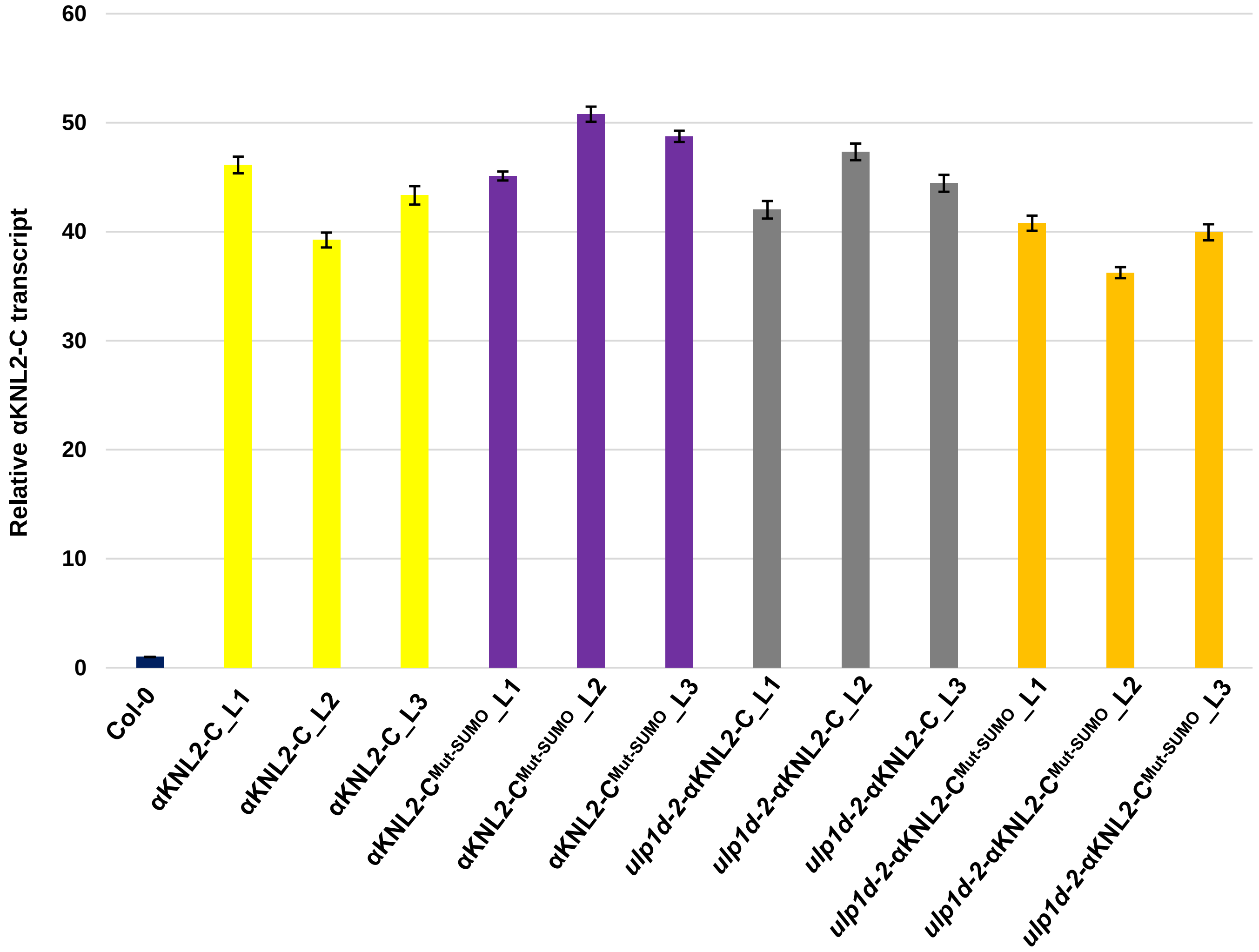


B

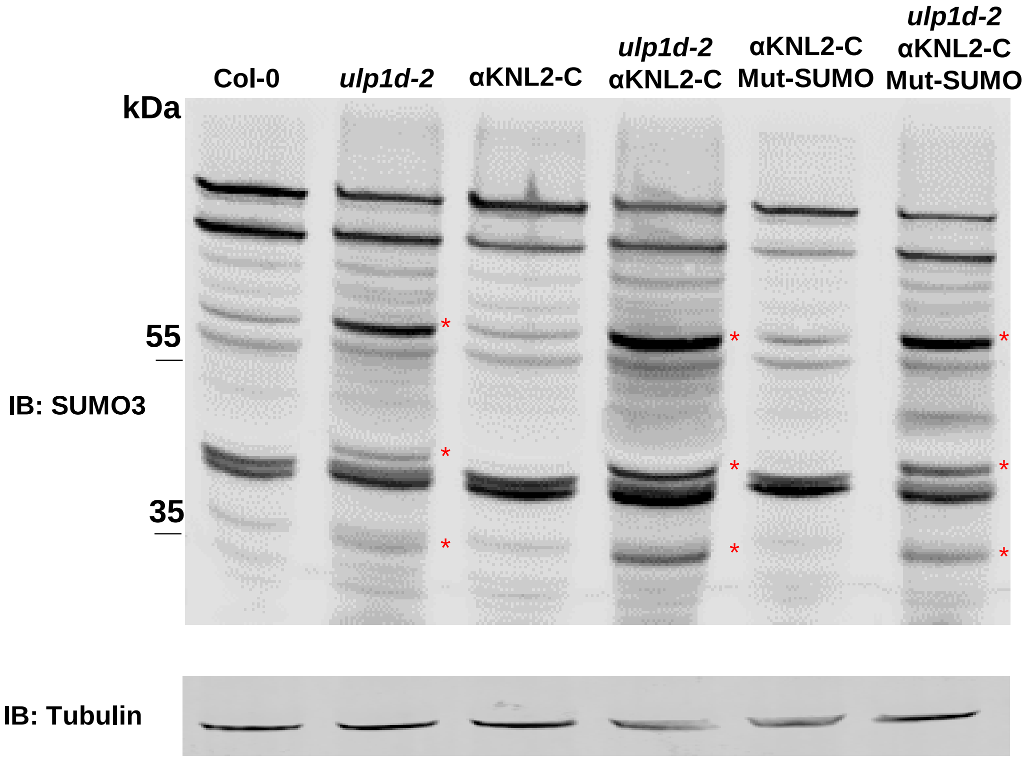


**Supplementary Figure 9. Transcript levels of *αKNL2* and SUMO3 western blot in αKNL2-C and αKNL2-C$^{Mut\text{-}SUMO}$ lines in wild-type and *ulp1d-2* mutant plants**

**(A)** Quantitative real-time PCR (RT-qPCR) analysis of αKNL2 transcripts in αKNL2-C and SUMOylation-deficient αKNL2-C$^{Mut\text{-}SUMO}$ independent Arabidopsis transgenic lines. Wild-type (Col-0) were included as controls. Transcript levels were normalized to *ACTIN2 and UBQ* expression. Similar transcript levels were detected across αKNL2-C and αKNL2-C$^{Mut\text{-}SUMO}$ transgenic lines in Col-0 and *ulp1d-2.* Data represent as mean ± SEM. Statistical analysis by ANOVA revealed no significant differences between the lines ($p > 0.5$). **(B)** Western blot analysis against anti-SUMO3 in the total protein extracts from wild-type, *ulp1d-2,* αKNL2-C, and αKNL2-C$^{Mut\text{-}SUMO}$ in wild-type and *ulp1d-2* mutants. The red asterisks shows the increase in the band intensities in *ulp1d-2* mutant background compared to wild-type (Col-0). Abbreviations: IB, Immunoblot.

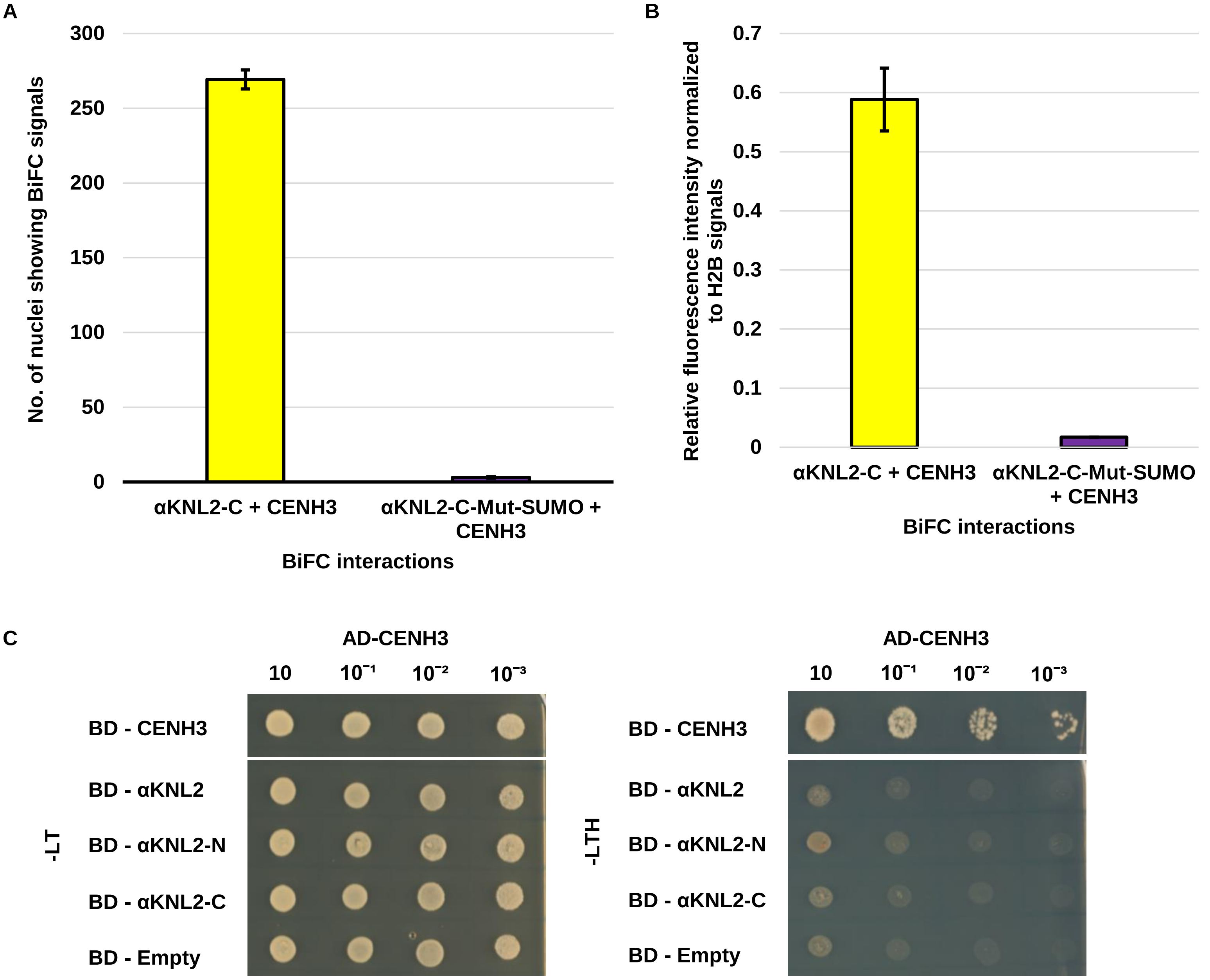


**Supplemental Figure 10. BiFC quantification and yeast two-hybrid assay for interactions between αKNL2 and CENH3** **(A)** Bar graphs represent the number of nuclei showing BiFC signals for αKNL2, αKNL2$^{Mut\text{-}SUMO}$ and CENH3 interactions. The number of nuclei showing BiFC signals was measured in $80 mm^2$ area. Data are presented as mean ± SEM. **(B)** The fluorescence intensity for BiFC signals were measured after normalization with H2B signals from 30 nuclei per sample (n = 30). Data are presented as mean ± SEM. **(C)** Zygotes expressing both prey CENH3 and bait (CENH3, αKNL2, αKNL2-N, αKNL2-C) are selected on -LT (Double dropout: YNB without Leu and Trp). Protein-protein interactions are assessed on -LTH (Triple dropout: YNB without Leu, Trp, and His). The strength of the protein-protein interactions was evaluated by a drop dilution test. AD, activating domain; BD, binding domain.

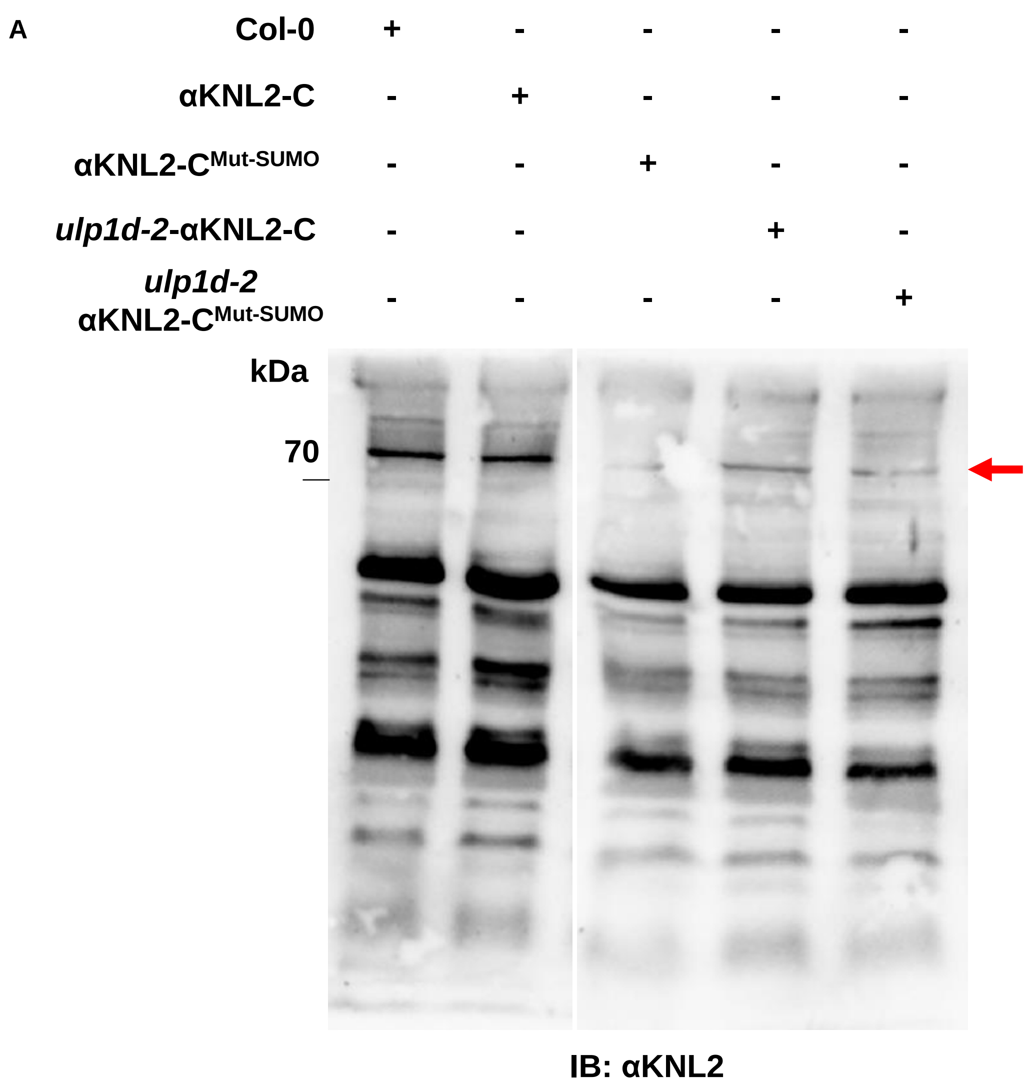


**Supplementary Figure 11. αKNL2 protein levels are reduced in the SUMOylation-deficient αKNL2-C mutant**

**(A)** Immunoblot analysis of αKNL2 in wild-type (Col-0), αKNL2-C-EYFP, and αKNL2-C$^{Mut\text{-}SUMO}$-EYFP in Col-0 and *ulp1d-2* background lines. The nuclear protein extracts were separated by SDS-PAGE and probed with anti-αKNL2 antibodies to detect endogenous αKNL2. The red arrow indicates the αKNL2 specific band. The tubulin control is same as Fig. 7C.
