## Supplementary Table for "The C-terminal SUMOylation-dependent regulation of αKNL2 governs its centromere targeting and interaction with CENH3"

**Supplementary Table 1. Primers used in this study**

| S. No | Gene | Forward primer | Reverse primer |
|---|---|---|---|
| Amplification of genes using attB primers | | | |
| 1 | SUMO1 (AT4G26840) | GGGGACAAGTTTGTACAAAAAAGCAGGCTTCATGTCTGCAAACCAGGAGG | GGGGACCACTTTGTACAAGAAAGCTGGGTGGCCGTAGCACCACCACCGC |
| 2 | SUMO2 (AT5G55160) | GGGGACAAGTTTGTACAAAAAAGCAGGCTTCATGTCTGCTACTCCGGAAGA | GGGGACCACTTTGTACAAGAAAGCTGGGTAAAGCAGAAGAGCTTCAGGC |
| 3 | SUMO3 (AT5G55170) | GGGGACAAGTTTGTACAAAAAAGCAGGCTTCATGTCTAACCCTCAAGATGA | GGGGACCACTTTGTACAAGAAAGCTGGGTAAGCCCATTATGATCGAAAAG |
| 4 | SUMO5 (AT2G32765) | GGGGACAAGTTTGTACAAAAAAGCAGGCTTCATGGTGAGTTCCACAGACAC | GGGGACCACTTTGTACAAGAAAGCTGGGTAGGAGTGTAAGGACCGCCACC |
| 5 | ULP1d (AT1G60220) | GGGGACAAGTTTGTACAAAAAAGCAGGCTTCATGACGAAGAGGAAGAAGG | GGGGACCACTTTGTACAAGAAAGCTGGGTTTACTCTGTCTGGTCACTGAC |
| 6 | ULP1d-N (AT1G60220) | GGGGACAAGTTTGTACAAAAAAGCAGGCTTCATGACGAAGAGGAAGAAGG | GGGGACCACTTTGTACAAGAAAGCTGGGTCTTACGGCGCCTTGAACTTTG |
| 7 | ULP1d-C (AT1G60220) | GGGGACAAGTTTGTACAAAAAAGCAGGCTTCATGAAATCAGAGGACACAGTG | GGGGACCACTTTGTACAAGAAAGCTGGGTCTCTGTCTGGTCACTGACACG |
| Primers used to confirm the positive entry and destination clones | | | |
| 8 | attB1 | GGGGACAAGTTTGTACAAAAAAGCAGGCTTC | |
| 9 | attB2 | GGGGACCACTTTGTACAAGAAAGCTGGGTC | |
| Primers used for PCR-based site-directed mutagenesis | | | |
| 10 | K378R-αKNL2C | AAACAAAAGGAGAATCGATGCGAG | TCCGCACTTTTGACTTTCGTCCCAG |
| 11 | K474R-αKNL2C | GAAAATCAAAGAGAAGTGAGAAGA | CTTTCGACAGGGGATCTTGAAATGC |
| 12 | K511R-αKNL2C | AATAAAGAGGAGAATCGACTTTG | TTTTCCCATGACAAGTTTTCTTCAG |
| 13 | Δ547-551-αKNL2C | CTAGAGTTTTGGCGTAACCAAATTC | CCTTCCTGATCTTGACCGTTTCTGT |
| 14 | Δ568-572-αKNL2C | GATGGTAGTGAGACTAACTCCGCTC | GTTCCGATCCATATCATAAACAGG |
| Primers used for cloning constructs for in vitro SUMOylation assay | | | |
| 15 | αKNL2 C-terminus (WT and SUMO mutant) to pET-Duet ; JJ225, JJ226 | ACCATCATCACCACAGCCAGATGAATTACTCTGGGACG | CTGAAAATACAGGTTTTCCGCTTTGATTTTCAAGTTTCTTCG |
| 16 | NSE2 to pET28 c+ ; JJ200, JJ201 | GGTGGACAGCAAATGGGTCGGATCCCCATGGCGTCGGCGTCCTCG | GGTGGTGGTGGTGGTGCTCGAGCTAATCTTCATCCACATCTTCTGTGAA |
| **Primers used for RT-qPCR analysis** | | | |
| 17 | qαKNL2-C | TCGACTTTGATGTGGAGGTAACAC | GAATCAGTAGACGCCGCATTGG |
| 18 | qCENH3 | GCAGGTCCAACTACGACCC | GCTGGTGAAGTTGTAGGATTTGT |
